# The spatially resolved tumor microenvironment predicts treatment outcome in relapsed/refractory Hodgkin lymphoma

**DOI:** 10.1101/2023.05.19.541331

**Authors:** Tomohiro Aoki, Aixiang Jiang, Alexander Xu, Yifan Yin, Alicia Gamboa, Katy Milne, Katsuyoshi Takata, Tomoko Miyata-Takata, Shaocheng Wu, Mary Warren, Celia Strong, Talia Goodyear, Kayleigh Morris, Lauren C. Chong, Monirath Hav, Anthony R. Colombo, Adele Telenius, Merrill Boyle, Susana Ben-Neriah, Maryse Power, Alina S. Gerrie, Andrew P. Weng, Aly Karsan, Andrew Roth, Pedro Farinha, David W. Scott, Kerry J. Savage, Brad H. Nelson, Akil Merchant, Christian Steidl

## Abstract

**PURPOSE:** About a third of relapsed or refractory classic Hodgkin lymphoma (r/r CHL) patients succumb to their disease after high-dose chemotherapy followed by autologous stem cell transplantation (HDC/ASCT). Here, we aimed to describe spatially resolved tumor microenvironment (TME) ecosystems to establish novel biomarkers associated with treatment failure in r/r CHL.

**METHODS:** We performed imaging mass cytometry (IMC) on 169 paired primary diagnostic and relapse biopsies using a marker panel specific for CHL biology. For each cell type in the TME, we calculated a ’spatial score’ measuring the distance of nearest neighbor cells to the malignant Hodgkin Reed Sternberg cells within close interaction range. ‘Spatial scores’ were used as features in prognostic model development for post-ASCT outcomes.

**RESULTS:** Highly multiplexed IMC data revealed shared TME patterns in paired diagnostic and early relapse/refractory CHL samples, whereas TME patterns were more divergent in pairs of diagnostic and late relapse samples. Integrated analysis of IMC and single cell RNA sequencing data identified unique architecture defined by CXCR5^+^ HRS cells and their strong spatial relationship with CXCL13+ macrophages in the TME. We developed a prognostic assay (‘RHL4S’) using four spatially resolved parameters, CXCR5+ HRS cells, PD1+CD4+ T cells, tumor-associated macrophages, and CXCR5+ B cells, which effectively separated patients into high-risk vs low-risk groups with significantly different post-ASCT outcomes. The RHL4S assay was validated in an independent r/r CHL cohort using a multicolor immunofluorescence assay.

**CONCLUSIONS:** We identified the interaction of CXCR5+ HRS cells with ligand-expressing CXCL13+ macrophages as a prominent crosstalk axis in relapsed CHL. Harnessing this TME biology, we developed a novel prognostic model applicable to r/r CHL biopsies, RHL4S, opening new avenues for spatial biomarker development.

## Introduction

Microenvironment biology has been extensively explored in various cancers, including lymphomas, and a large number of studies have revealed the pathologic importance of reactive immune cells in the tumor-microenvironment (TME)^1–3^. Classic Hodgkin Lymphoma (CHL) is unique amongst virtually all cancers as the malignant Hodgkin and Reed Sternberg (HRS) cells are greatly outnumbered by reactive, non-neoplastic cells in the TME^4, 5^. Typically, the malignant HRS cells represent less than 1% of cells in an individual tumour. Despite progress in elucidating TME biology and the extensive cellular crosstalk by cytokines/chemokines as a hallmark feature of CHL pathogenesis^6, 7^, the molecular determinants of treatment failure remain mostly unknown.

Despite recent treatment advances, about a third of relapsed and refractory (r/r) CHL patients succumb to their disease after high-dose chemotherapy followed by autologous stem cell transplantation (HDC/ASCT)^8, 9^. Recent studies have reported that gene expression signatures representing non-neoplastic cells of the TME are associated with outcomes following therapy^7, 10, 11^. Our group previously developed and validated a clinically applicable prognostic assay (RHL30), which identifies a subset of patients at high-risk of treatment failure following HDC/ASCT in r/r CHL^12^. The study also established that relapse biopsies are superior to diagnostic biopsies to develop biomarker assays for predicting post-ASCT outcomes. However, RHL30 does not incorporate important information about cellular interactions, specific expression features of HRS cells, and the spatial architecture of the TME. Recent progress with multiplex imaging techniques has enabled comprehensive spatial characterization of TME biology, suitable for formalin-fixed paraffin-embedded tissue (FFPET). This was demonstrated in our recent publication using imaging mass cytometry (IMC) in primary CHL where novel interactions between tumor-specific immunosuppressive cell populations and malignant HRS cells were described^13^. We therefore hypothesized that the more detailed description of spatially resolved ecosystems in samples at the timepoint of clinical relapse would lead to the development of refined biomarkers with the potential to improve risk stratification for post-secondary treatment outcomes.

Towards this goal, we performed IMC analysis of paired pre-treatment/relapse biopsies (n=142) and non-relapse biopsies (n=22) to describe the unique changes in microenvironment architecture linked to relapse on a per-patient basis. Our data shed light on the unique interactions between cancer cells and the TME, highlighting the spatial interaction between CXCR5+ HRS cells with CXCL13+ macrophages as a characteristic feature of r/r CHL. Consequently, we developed a novel spatially resolved prognostic biomarker assay, “RHL4S”, that was also translated into a multi-color immunofluorescence (MC-IF) assay suitable for future implementation in routine pathological workflows.

### PATIENTS AND METHODS

Detailed materials and methods are available in the Supplementary Data file.

#### Patient Cohort

We analyzed IMC data from 165 CHL samples, including 71 patients with paired primary diagnostic and relapse specimens and 22 diagnostic control samples without relapse, termed the ‘discovery cohort’ (**Appendix Fig A1)**^12^. Biopsies of these CHL cases, treated at BC Cancer between 1985 and 2011, are part of a tissue microarray (TMA) that was previously reported^12^. In brief, the patients were selected according to the following criteria: received first-line treatment with doxorubicin, bleomycin, vinblastine, and dacarbazine (ABVD) or ABVD-equivalent therapy with curative intent; experienced CHL progression/relapse after first-line treatment and tissue derived from an excisional biopsy was available^12^. Patients were classified as having early relapse if their CHL progressed within 12 months after initial diagnosis or was refractory to first-line treatment. We also assembled a second, independent cohort for validation, termed the ‘validation cohort’ (n = 44) used for MC-IF. For the validation cohort, we selected r/r CHL patients treated at BC Cancer between 2012 to 2021 with available FFPET relapse biopsies according to the same selection criteria as the discovery cohort. Patient characteristics of the validation cohort are summarized in **Appendix Table 1 and Appendix FigA1**.

This study was reviewed and approved by the University of British Columbia-BC Cancer Agency Research Ethics Board (H14-02304), in accordance with the Declaration of Helsinki.

#### Survival analysis

Overall survival (OS) was defined as time from CHL diagnosis to death from any cause. Time to first relapse was defined as time from primary diagnosis to first CHL progression/relapse, or death from CHL. Post-ASCT-OS was defined as time from ASCT to death from any cause. Post-ASCT failure free survival (FFS) was defined as time from ASCT to CHL progression/relapse, or death from any cause. Non-parametric survival analyses with a single binary predictor were analyzed using the Kaplan-Meier method and results were compared using the log rank test. Univariate and multivariate Cox regression analyses were performed to assess the effects of prognostic factors. Survival analyses were performed in the R statistical environment (v4.2.2).

#### Imaging mass cytometry

IMC was performed using a customized marker panel for CHL on 5µm sections of the TMAs containing 1.5mm duplicate cores of each sample of the discovery cohort **(Appendix Table A2)**. Slides were imaged using the Fluidigm Hyperion IMC system with a 1µm laser ablation spot size and frequency of 200Hz. Tissue areas of approximately 1 mm^2^ per sample were ablated and imaged. Segmentation was performed by the Ilastik machine learning package^14^ in conjunction with the CellProfiler IMC Segmentation Pipeline^15^. We used the Phenograph clustering algorithm to identify major cell types and functional cellular subsets and defined each immune cell population based on marker expression. For each cell type, we calculated a ‘spatial score’, defined as the term: (1 - average distance of HRS cells to the 5 nearest neighbor cells of that type capped at 50 micrometers) to distinguish relationships between HRS cells and clusters of interacting cells. This strategy enabled us to quantify the spatially resolved cellular architecture in CHL (**Appendix Fig A2**, See appendix for the detail)^16^.

#### Prognostic model development

To develop a prognostic model for r/r CHL, we applied our new cross-format LASSO (Least Absolute Shrinkage and Selection Operator) plus algorithm on the combination of two separate lists of standardized ‘spatial scores’ and standardized traditional protein-based cellular abundance percentages. This analysis enabled us to select six variables associated with post-ASCT FFS with p-values <= 0.05. To improve the practicality of the model, we further removed two variables whose 95% confidence intervals (CI) overlapped with the range of 0.95-1.05. Using the spatial scores of these four selected variables, we developed a risk classification prediction model, RHL4S based on our new XGpred algorithm in cvmpv R package (https://github.com/ajiangsfu/csmpv), which distinguished high and low-risk groups of patients with respect to post-ASCT FFS using an optimal threshold (see Appendix for details).

#### Prognostic model translation using MC-IF

MC-IF was performed as previously described^13^. In brief, TMA slides were deparaffinized and incubated with two panels of six antibodies to each marker of interest **(Appendix Table A3)**, followed by detection using Mach2 HRP and visualization using Opal fluorophores. To analyze the spectra for all fluorophores included, inForm image analysis software (v2.4.10; PerkinElmer, USA) was used (see Appendix for details).

To establish a prognostic model that can be generalized and is applicable to routine pathology procedures, we built a simplified panel which contains the multiparametric and spatial information of the 4 variables from the IMC-based model (**Appendix TableA3**). Spatial scores were derived from the MC-IF data and scores of the risk classification model were calculated per patient. To confirm the technical concordance between spatial scores derived from IMC vs MC-IF, we applied the MC-IF panel to a subset of the discovery cohort (N = 19). To adjust RHL4S scores between IMC and MC-IF methodologies, calibration was performed correcting scores by a calibration value defined as the mean difference between RHL4S IMC and MC-IF scores **(Appendix Fig A3-4)**.

#### Single cell RNA sequencing and library preparation

Samples were processed and libraries were prepared for single cell RNA sequencing (scRNA-seq) as previously described^13^. In brief, sorted cells on a FACS Fusion (BD Biosciences) using an 130 µm nozzle (**Appendix FigA5 and Appendix Table A4**) from cell suspensions were collected^17, 18^, and enriched HRS cells were loaded into a Chromium Single Cell 5’ Chip kit v2 (PN-120236). Libraries were constructed using the Single 5’ Library and Gel Bead Kit v2 (PN-120237) and Chromium i7 Mulitiplex Kit v2 (PN-120236). The library was measured using Agilent Bioanalyzer High Sensitivity chip and Qubit dsDNA HS Assay Kit and run on Illumina Nextseq550. To boost transcriptome information for cell-to-cell interaction analyses, hybridization capture of marker genes (**Appendix Table A5**) was performed as described previously^19^ using Twist Hybridization and Wash Kit (Twist Bioscience). The captured library was measured using Agilent Bioanalyzer High Sensitivity chip and Qubit dsDNA HS Assay Kit and run on Illumina Nextseq550 (**see Appendix for details**).

## Results

### The spatially resolved TME of r/r CHL

Using a customized marker panel specific for CHL biology (**Appendix Table A2**), we obtained highly multiplexed images for a total of 7,146,042 cells at 1 um^2^ resolution. To define the spatial architecture and related cellular interactions in r/r CHL, we first investigated the differences in TME characteristics according to disease status (initial diagnosis *vs* early relapse *vs* late relapse), showing differential abundance of each immune cell type (**Fig. 1A**). Intriguingly, while diagnostic samples demonstrated known CD4 T cell enrichment patterns, relapse samples showed a unique immune cell abundance profile. Since high-resolution characterization beyond cellular abundance is needed to delineate the complexity of the cellular ecosystem in the TME^20^, we defined comprehensive TME ecotypes using unsupervised clustering by leveraging a multitude of TME features (**Appendix, Fig. A6).** When determining TME ecotypes in individual diagnostic and relapsed biopsy pairs per patient, we observed significant differences in TME dynamics between early relapse and late relapse samples, with a significantly increased number of TME type transitions in late relapse (p < 0.01) (**Fig 1B**). Late relapse samples were characterized by an abundance of non-malignant B cells (P < 0.01) along with CD4 (P < 0.05) or CD8 T cell enrichment (P < 0.01) (**Fig. 1A, C and D).** Conversely, CHL biopsies from patients with early relapse demonstrated more similar TME patterns between diagnostic and relapse samples (**Fig 1B**). In particular, the macrophage/myeloid cell enriched TME ecotypes 1 and 6 were constant over time between diagnostic and early relapse biopsies (**Fig 1B and C**). Further analyses revealed that a CD163+ macrophage population^21^, indicative of M2 polarization, was significantly enriched in early relapse samples (**Fig. 1D and E**). Consistent with these findings, spatial analyses of relapse samples revealed an inverse correlation of CD163+ macrophages and B cells in cellular neighborhoods (**Fig. 1F**).

**Figure 1.**
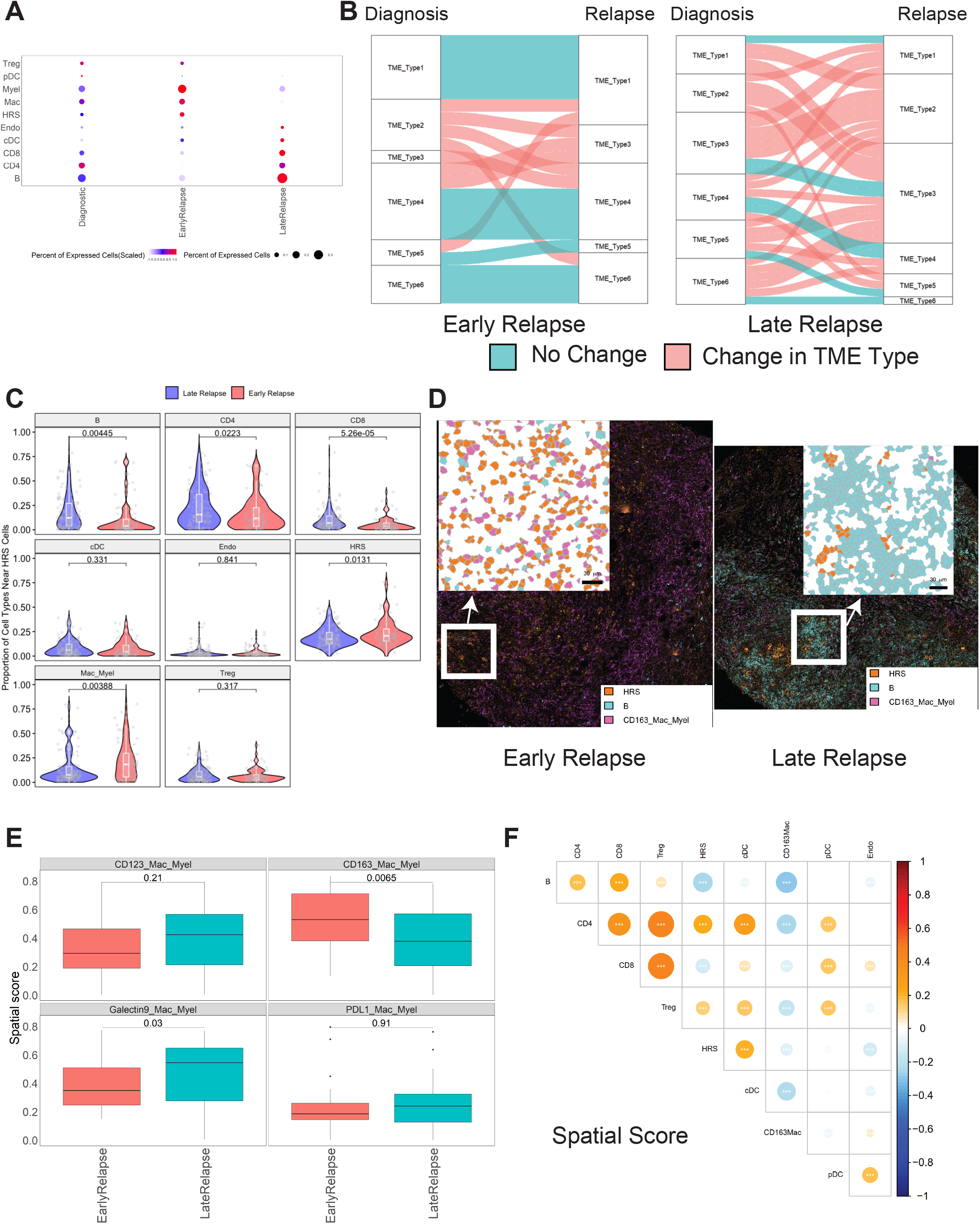
Distinct spatially resolved tumor microenvironment features according to relapse status. **A.** Proportion for the indicated immune cell population by imaging mass cytometry (IMC)-based cluster assignment datasets. **B.** The alluvial plot shows the tumor-microenvironment (TME) types and their dynamic change between diagnostic samples and relapse samples according to relapse status. Horizontal ribbons represent individual cases and can be followed from left to right. Blue color of the ribbons indicates that there is no TME type change between diagnostic and relapse samples while red colored samples indicate change of TME type. **C.** Violin plot indicating the spatial score for the indicated cell types near Hodgkin and Reed Sternberg (HRS) cells according to relapse status. **D.** IMC analysis from FFPE sections of classic Hodgkin lymphoma (CHL) shows localization of immune cells according to relapse status. A representative case with early relapse CHL case (left) shows numerous CD163+ macrophage/myeloid cells and rare B cells. In contrast, a representative late relapse HL case (right) shows few CD163+ macrophage/myeloid cells and abundant B cells. **E.** Box plot indicating the spatial scores of macrophage/myeloid cell subtypes near Hodgkin and Reed Sternberg (HRS) cells according to relapse status. **F**. Dot plot showing correlation of spatial scores of major immune cell markers by imaging mass cytometry (IMC). Dot size and color summarize Pearson correlation values, with positive correlations represented in red and negative correlations represented in blue. Asterisks represent associated p-values (*P < 0.05; **P < 0.01; ***P < 0.001).

### The unique spatial architecture associated with CXCR5^+^ HRS cells in relapse biopsies

In previous studies, the most prominent obstacle for biomarker development in CHL was the scarcity of the malignant HRS cells and the heterogeneity of TME composition within individual tumor biopsies. Our IMC panel was designed to simultaneously quantify protein expression on HRS cells and the TME, including known variably expressed markers on HRS cells, such as CD30, PD-L1 and major histocompatibility class I and II (MHC-I and MHC-II). Unsupervised clustering using the phenograph tool identified several new subsets within the HRS cellular compartment, defined by, for example, high GATA3 and CXCR5 expression. These phenotypic HRS cell definitions are additive to other known subsets, such as HRS cells with high PD-L1 or CD123 expression^22, 23^ (**Appendix Fig A7**). Next, we evaluated the prognostic impact of these HRS features in the spatial context of r/r CHL. In particular, LASSO analysis identified CXCR5+ HRS cells as the most significant HRS cell phenotype correlated with post-ASCT FFS, in contrast to the absence of an outcome correlate with all HRS cells, not further specified (**Fig. 2A, Appendix Fig A8**). To confirm CXCR5 expression patterns on HRS cells by both protein and RNA levels, we assessed expression of CXCR5 using immunohistochemistry (IHC) and reanalyzed published Affymetrix gene expression data generated from micro-dissected HRS cells of primary HL samples^24^. These analyses confirmed variable CXCR5 surface protein expression was well correlated with mRNA expression, and CXCR5 was highly expressed in a subset of CHL tumors (**Fig. 2B**) comparable to expression levels found in CD77+ germinal center B cells (**Fig 2C**). Interestingly, CXCR5+ HRS cells were spatially arranged together with other CXCR5+ HRS cells forming cell clusters as an architectural feature (P < 0.05). CXCR5+ HRS cell clusters were also characterized by a lower abundance of non-malignant immune cells, including CD4+ T reg cells and CD20+ B cells when compared to CXCR5-HRS cells (**Fig. 2D-F**), suggesting distinct TME characteristics associated with CXCR5+ HRS cells.

**Figure 2.**
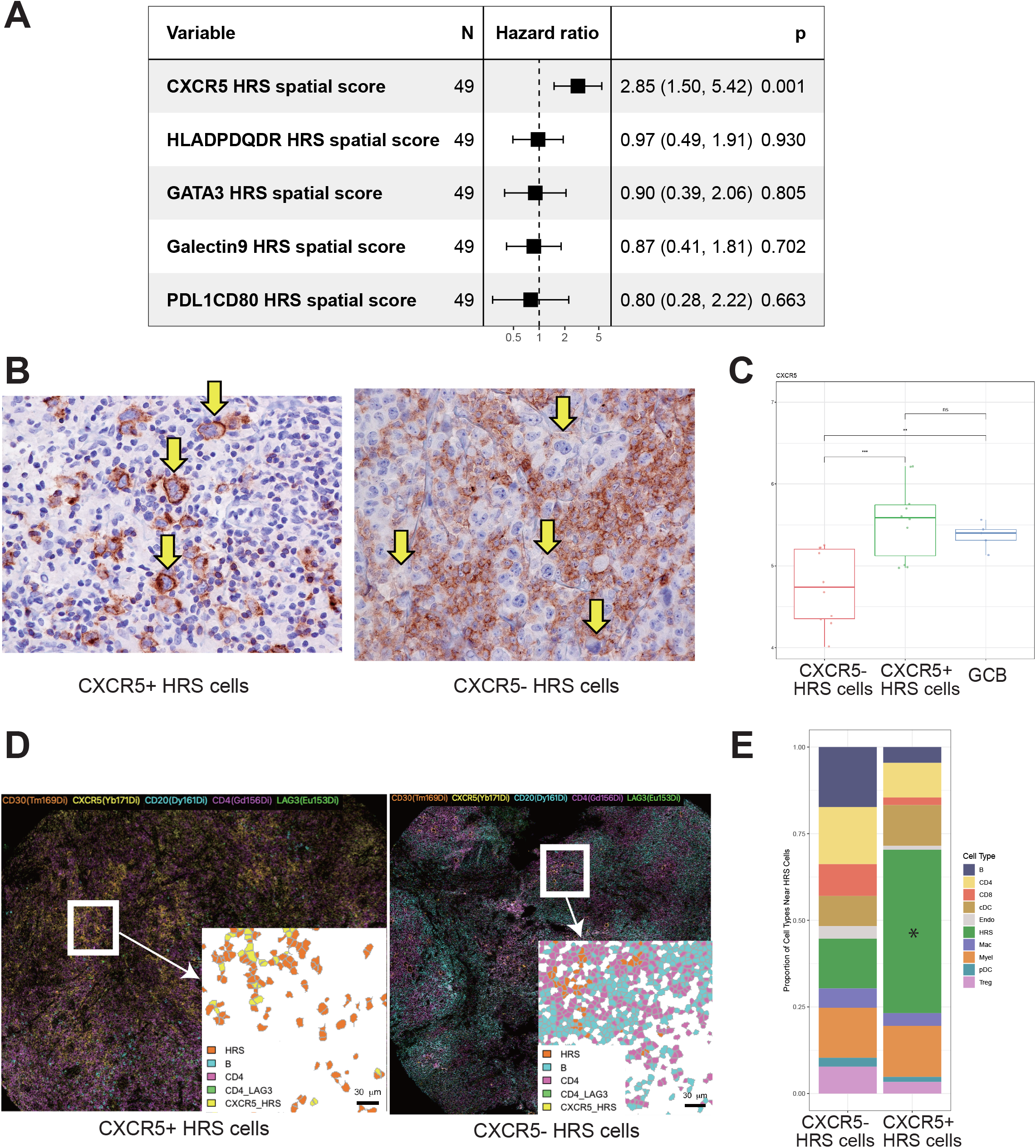
Characteristics of the tumor-microenvironment of classic Hodgkin lymphoma (CHL) associated with CXCR5 positivity on HRS cells. **A.** Forest plots summarize the prognostic factors in relapsed classic Hodgkin lymphoma treated with HDC/ASCT according to HRS cells features by imaging mass cytometry (IMC). **B**. IHC staining for CXCR5 in representative cases with either positive (Left) or negative (Right) HRS cells (×400). **C.** The expression of CXCR5 in microdissected HRS cells from primary CHL samples (separated by CXCR5 status evaluated by IHC) and germinal center cells from reactive tonsil tissue (GCB; t test; ns: P > 0.05; *, P ≤ 0.05; **, P ≤ 0.01). **D.** IMC image for selected immune subsets in representative cases with either CXCR5 positive (Left) or negative (Right) HRS cells. **E**. Relative proportion of cell subtypes near either negative (Left) or positive (Right) HRS cells. *P < 0.05.

We next sought to understand the cellular interactions between CXCR5^+^ HRS cells and other immune cell populations. Although we observed fewer TME components surrounding CXCR5^+^ HRS cells in the IMC data (**Fig. 2E**), we hypothesized that previously unknown immune cell populations, which cannot be defined with the current IMC panel, might interact with CXCR5^+^ HRS cells. Therefore, we additionally performed scRNA-seq on CHL samples with partial enrichment of the HRS cell population by cell sorting (**Appendix Fig A7**). Using the cell-to-cell communication tool, Cell Chat^25^, we predicted CXCR5-CXCL13 interaction between CXCR5^+^ HRS cells and CXCL13^+^ macrophages (**Fig. 3A, Appendix Fig A9)**. This significant interaction was also validated by the alternative iTALK method^26^ (**Fig. 3B**). CXCL13 is a well-described cell attractant via the CXCL13/CXCR5 axis. Intriguingly, CXCL13+ macrophages mostly (> 99%) did not co-express M2 macrophage markers, such as CD163 or CD206^11^ (**Appendix Fig A10**), indicating a distinct profile of this population. Since CXCL13 was not part of the original IMC panel, we were not able to study its expression in the IMC data. Therefore, to further describe the spatial relationship between CXCR5+ HRS cells and CXCL13+ macrophages, we applied multicolor immunofluorescence (MC-IF) on the discovery cohort TMA that was used for IMC. Consistent with the results of the scRNAseq-based cell-to-cell interaction prediction, we observed a strong positive correlation of the abundance of CXCR5+ HRS cells with CXCL13+ macrophages in the MC-IF data (**Fig. 3C)**. Furthermore, CXCL13+ macrophages were located in close proximity to CXCR5+ HRS cells (**Fig. 3D and 3E)**, supporting the importance of the CXCR5/CXCL13 axis in CHL.

**Figure 3.**
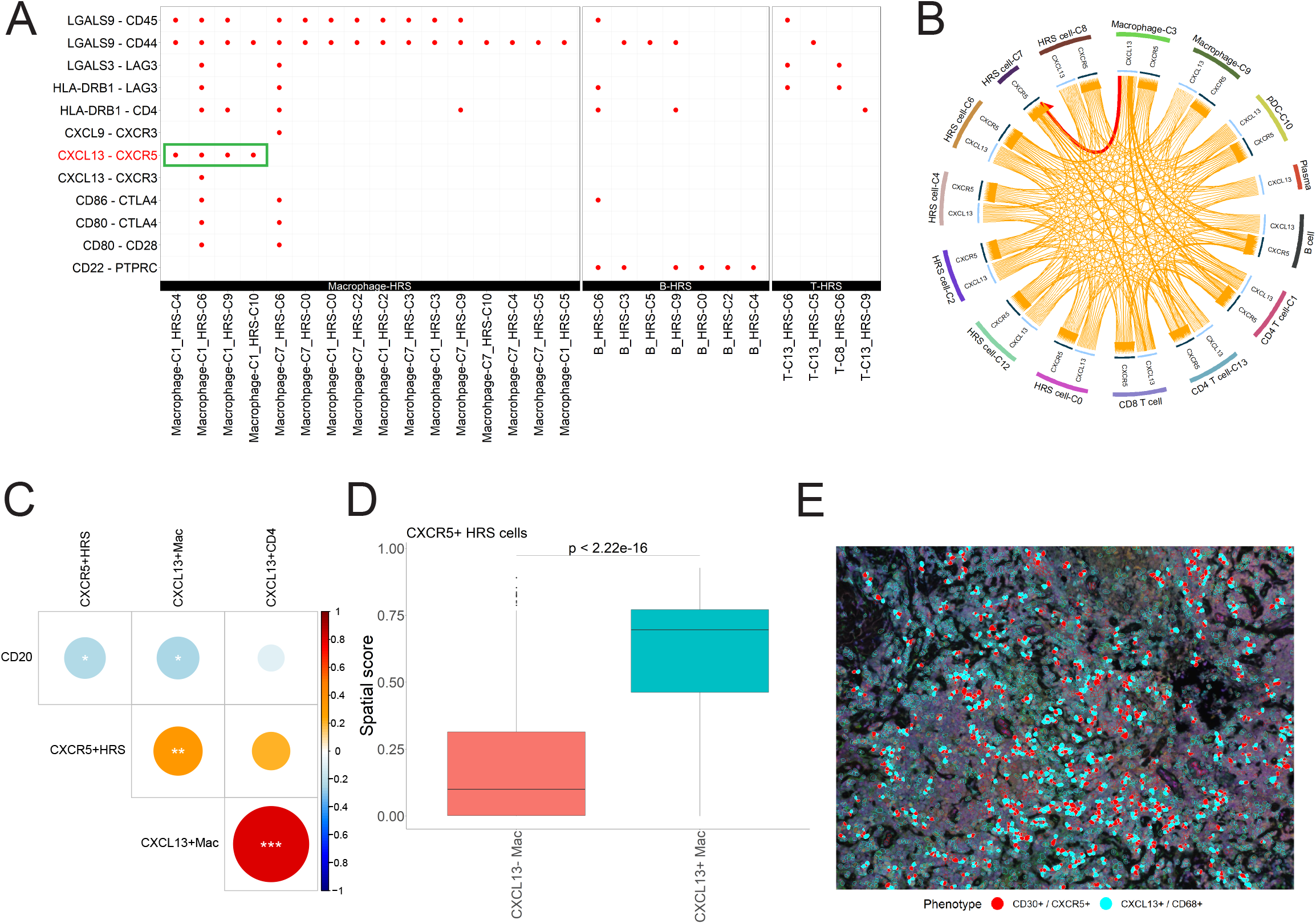
CXCL13/CXCR5 interaction in CHL. **A.** The dot plot shows significant ligand and receptor interaction between HRS cells and immune cell populations using Cell Chat. **B**. An interaction between CXCL13 and CXCR5 on immune cells and HRS cells in CHL samples was predicted using the iTALK tool. **C**. Dot plot showing correlations of the proportions of selected immune cell subsets with emphasis on CXCL13/CXCR5 interaction (MC-IHC). Dot size and color summarize Pearson correlation values, with positive correlations represented in blue and negative correlations represented in red. Asterisks represent associated p-values (*P < 0.05; **P < 0.01; ***P < 0.001). **D**. Boxplot showing the spatial score of CXCL13+ and CXCL13-macrophages in the region surrounding CD30+ cells (HRS). **E**. Membrane map depicting CD68+CXCL13+ macrophages (light blue) and CD30+CXCR5+ HRS cells (red).

### Development of a prognostic model leveraging spatial TME information

Next, we sought to construct a prognostic model for post-ASCT outcomes in r/r CHL patients. We anticipated that IMC analysis would take advantage of simultaneously capturing HRS cell and TME biology in directly surrounding regions. Considering the previous demonstration of the superiority of using relapse specimens for outcome prediction in r/r CHL^12^, we focused our analyses on biomarker measurements in relapse samples. For feature selection, we first performed cross-format LASSO plus analysis on two separate sets of variables, namely standardized spatial scores and conventional cellular abundance percentages (based on protein expression markers) (**See Method, Appendix file**). Strikingly, all top five variables, which were significantly associated with post-ASCT FFS p-values <= 0.05, were ’spatial scores’ (**Fig 4A)**, confirming the importance of spatially-informed parameters for outcome prediction. Consistent with the independent prognostic importance of these variables (**Fig. 4A**), each patient sample showed distinct spatial patterns which were linked to these cellular components (**Fig. 4B**). Four variables, CXCR5+ HRS cells (hazard ratio [HR] (95% CI): 2.86 (1.63-5.02)), PD1+ CD4+ T cells (HR (95% CI): 2.80 (1.46-5.35)), macrophages (HR (95% CI): 1.99 (1.14-3.47)) and CXCR5+ B cells (HR (95% CI): 0.19(0.06-0.57)) were identified as factors most significantly associated with post-ASCT FFS.

**Figure 4.**
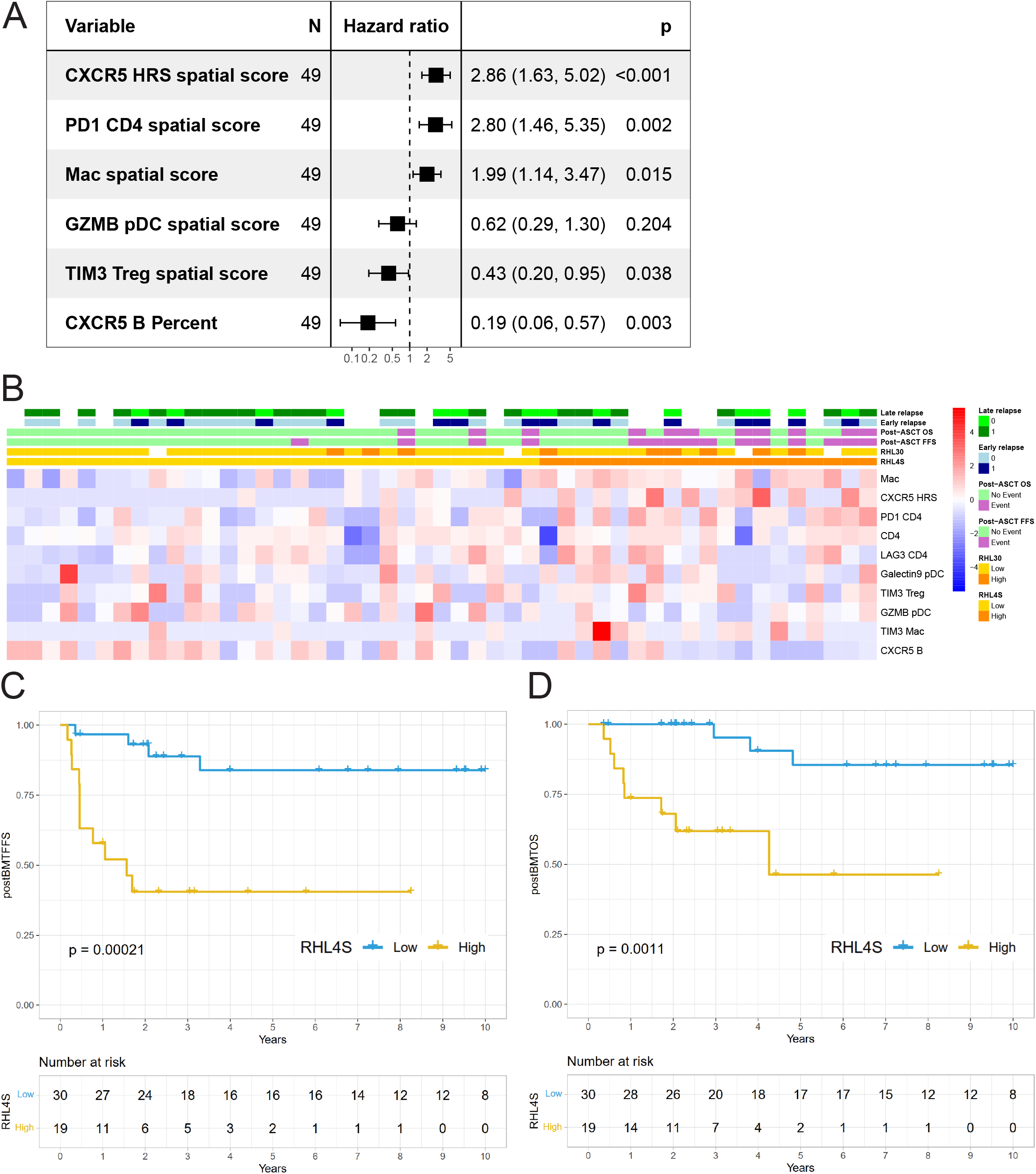
Development of a novel prognostic model, RHL4S, which predicts failure free survival after autologous stem cell transplantation (ASCT). **A.** Forest plots summarize the prognostic factors in relapsed classic Hodgkin lymphoma treated with HDC/ASCT according to imaging mass cytometry (IMC). **B**. Heatmap of the spatial scores in RHL4S according to IMC. Cases are ordered by RHL4S model score. Kaplan-Meier curves of the high-versus low-risk groups for (**C**) post-ASCT failure-free survival (FFS) and (**D**) post-ASCT overall survival (OS) as identified by RHL4S. P values were calculated using a log rank test.

To leverage the multifactorial spatial biologic features which are linked to relapsed/refractory biology, we developed a risk prediction model, RHL4S, using the spatial scores from these four variables. RHL4S identified a high and low-risk group of patients with the high-risk group of patients having significantly inferior post-ASCT FFS (5-year post-ASCT FFS: 41% vs 81%; P < .0001; **Fig 5A**) and inferior post-ASCT OS (5-year post-ASCT OS: high-risk, 46 % vs low-risk, 85%; P = .001; **Fig 5B**).

**Figure 5.**
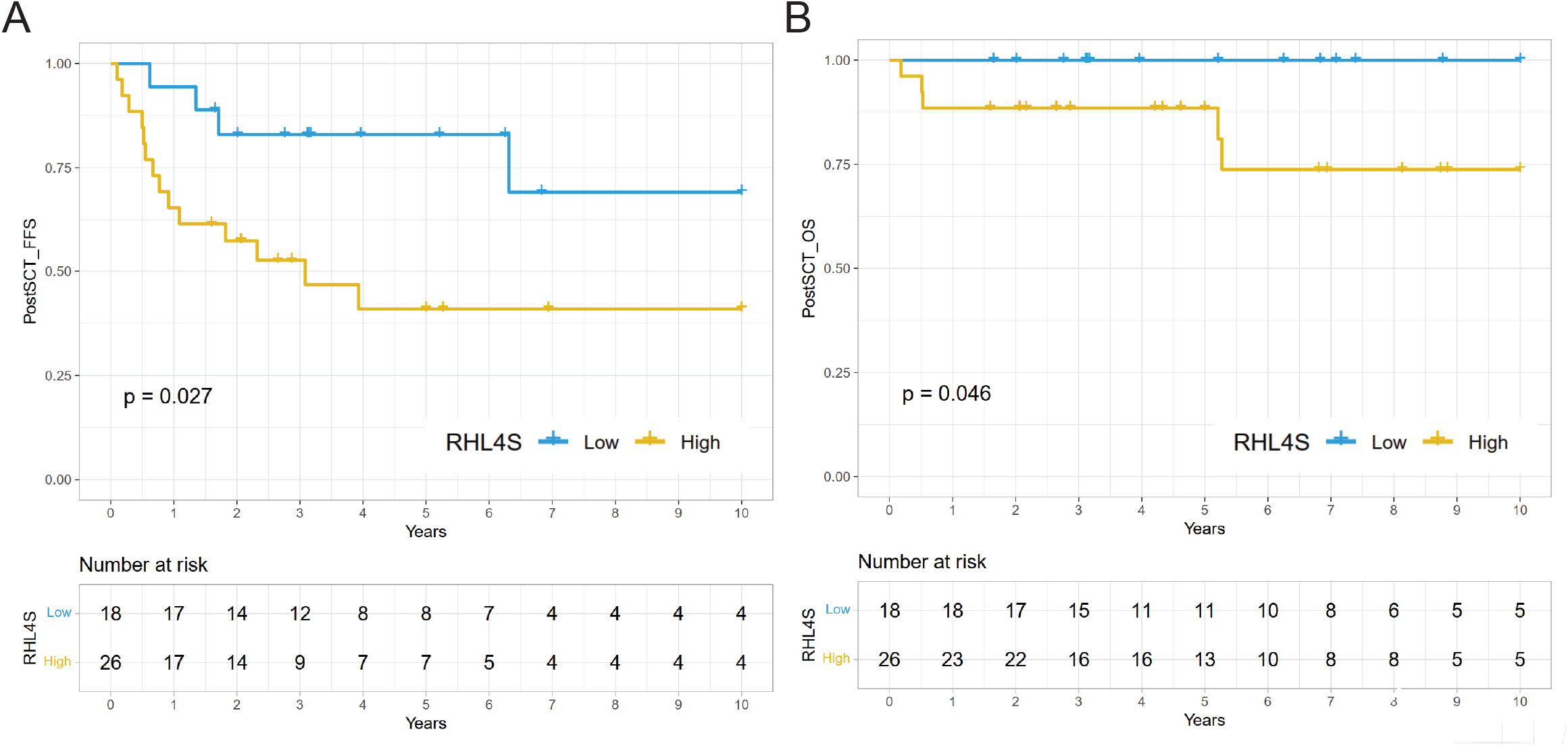
Validation of RHL4S in independent cohort of relapsed and refractory classic Hodgkin Lymphoma (r/r CHL). Kaplan-Meier curves of the high-versus low-risk groups for (**A**) post-ASCT failure-free survival (FFS) and (**B**) post-ASCT overall survival (OS) as identified by RHL4S in the independent validation cohort, respectively. P values were calculated using a log rank test.

We then investigated the independent prognostic value of RHL4S with respect to previously reported prognostic factors of post-ASCT outcomes including RHL30 (**Fig 4B**), time to first relapse, chemo-resistance, B symptoms at relapse, age ≥ 45 years at ASCT, and stage IV disease at initial diagnosis. Pairwise multivariable Cox regression analysis including RHL4S and these known prognostic variables demonstrated that the RHL4S risk group was statistically independent (P < .05) **(Appendix Fig 11)**.

### Independent Validation for RHL4S

Towards the goal of establishing a prognostic model that can be readily used for diagnostic procedures in clinical practice, we next translated the IMC-based RHL4S assay into MC-IF methodology. We built a simplified marker panel encompassing the four expression variables of the predictive model (**Appendix TableA3 and Appendix Fig A12**) and calculated calibrated individual spatial scores from the MC-IF data in the independent validation cohort of relapse biopsies from 44 r/r CHL patients uniformly treated with second-line chemotherapy and consolidating ASCT (**Table 1**). Consistent with the finding in the discovery cohort, high-risk patients displayed unfavorable post-ASCT FFS (5-year post-ASCT FFS: 41% vs 83%; P = .027; **Fig 5C**) and post-ASCT OS (5-year post-ASCT OS: 88% vs 100%; P = .046; **Fig 5D**) in the independent validation cohort. Finally, we investigated the independent prognostic value of RHL4S with respect to previously reported prognostic factors of post-ASCT outcomes in the validation cohort. Pairwise multivariable Cox regression analysis demonstrated the independence of RHL4S (P < .05) or trends toward independence against these markers **(Appendix Fig A13)**. Of note, although the mRNA gene-expression-based RHL30 assay showed similar performance to RHL4S in the discovery cohort, RHL30 did not show a significant difference between the high and low-risk groups in the validation cohort **(Appendix Fig A14)**, indicating superior performance of RHL4S.

## Discussion

Here, we have developed and validated a novel prognostic model, RHL4S, based on spatial TME biology, that can predict outcome after ASCT in patients with r/r CHL. Our study establishes a paradigm for the strong prognostic value of spatially resolved biomarkers over traditional expression-based biomarkers in lymphoma and suggests the biological importance of tumor architectural patterns for biomarker development and discovery of immunotherapeutic targets in other cancer types. Importantly, RHL4S was associated with post-ASCT outcomes regardless of the relapse status (early relapse or late relapse).

The main goal of treatment in CHL is the achievement of high cure rates while maintaining minimal toxicity. With recent advances of targeted treatments such as the anti-CD30 antibody drug conjugate brentuximab vedotin (BV), and PD-1 blockade (nivolumab and pembrolizumab) in r/r CHL^27–30^, the establishment of reliable risk-stratification at the time point of relapse, to discriminate patients with favorable vs unfavorable post-secondary treatment outcomes, has emerged as a priority. Ideally, risk stratification should be achieved before or during the early phase of second-line treatment to adapt treatment modality and intensity and to complement response-adapted approaches such as interim-PET and dynamic tumor burden markers (e.g. TARC or ctDNA measurements). While the gene-expression based model, RHL30, has been previously developed towards the same goals^10^, the clinical utility of this gene expression-based assay as a predictive biomarker is limited due to problems with routine accessibility of digital gene expression assays in pathology labs. By contrast, RHL4S can be implemented on technology platforms of MC-IF, which can be more easily implemented in routine diagnostics under high quality standards such as the Clinical Laboratory Improvement Amendments program^31^.

RHL4S includes four spatially resolved variables, including macrophages that were identified as a prognostic biomarker in CHL based on raw cellular abundance in the TME in multiple studies^7, 32–34^. We now also identified a phenotypically defined subset of HRS cells (CD30+CXCR5+) that was associated with outcome, and intriguingly, we observed enrichment of CXCL13+ macrophages in regions surrounding CXCR5+ HRS cells. We also recently found that CXCL13+PD1+ TFH-like cells are enriched in a specific subtype of CHL (lymphocyte-rich CHL) and associated with poor clinical outcome in this rare subtype^6^. These two studies indicate the importance of ligand (CXCL13) and receptor (CXCR5) interaction and their association with treatment failure in CHL. Importantly, CXCL13/CXCR5 targeting agents are currently under investigation in hematological malignancies and in autoimmune disease^35–37^. Therefore, additional investigations into the CXCR5/CXCL13 biology are warranted to determine therapeutic benefit in CHL. Moving forward, the impact of RHL4S on CHL patients treated with immunotherapy before HDC/ASCT needs to be determined as the promising efficacy of PD-1 blocking antibodies in the first/second line setting has changed the overall view on treatment strategies in CHL^9, 38^.

Our study also highlights the contrast between early relapse and late relapse biology. We observed relatively similar TME features between diagnostic and relapse biopsies in early relapse cases, while differences were more pronounced in late relapse cases. This finding raises the hypothesis that the establishment of malignant cellular ecosystems, including HRS cell-driven shaping of the TME, is persistent over time after first-line treatment in early relapse CHL, and by contrast, the biology of late relapses is indicative of more divergent disease and more dynamic changes in the TME. The latter scenario might also be supported by reports of clonally independent malignant HRS cells between diagnostic and relapse time points^39^.

The multiple cellular components associated with treatment outcome strongly suggest that the mechanism underlying relapsed disease might not be uniform (**Fig 6**). Nevertheless, delineating specific and targetable biology underlying relapsed/refractory disease, such as the CXCR5-CXCL13 axis, might lead to more effective delivery of precision oncology. The development of biology-driven biomarkers in CHL might also help pave the way for rational treatment selection, an approach that is currently emerging in diffuse large B-cell lymphoma based on gene expression and mutational profiling^40–42^. While RHL4S shows promise in guiding treatment selection and might compliment response-adapted treatment algorithms, further studies are needed to investigate the clinical utility of RHL4S in patients treated with immunotherapy-based treatment and potential transplantation-free treatment strategies in the future. Moving forward, we anticipate that the value of spatially resolved biomarkers will be tested in additional lymphoma subtypes^43^ and other cancers^44^ with a biologically important TME component.

**Figure 6.**
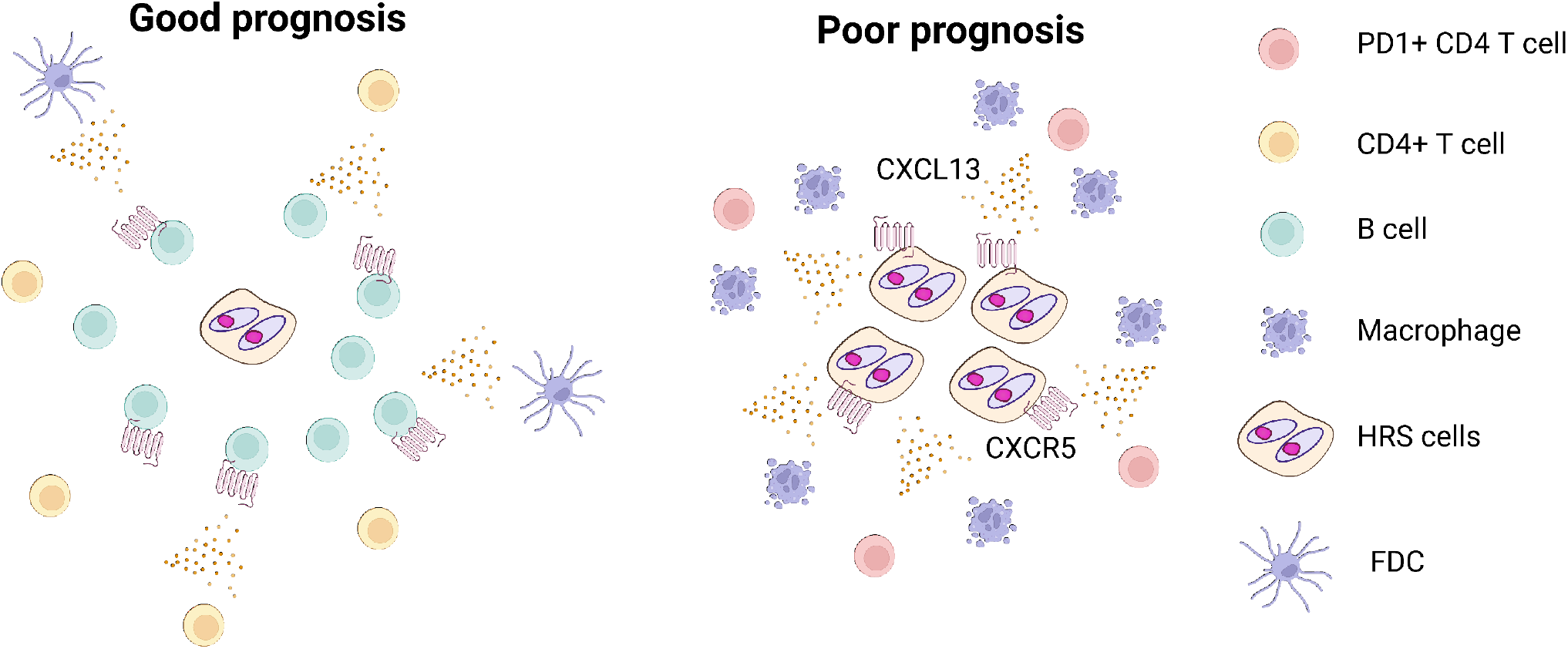
Graphical summary of findings in relapsed and refractory classic Hodgkin Lymphoma (r/r CHL). r/r CHL with poor prognosis is characterised by CXCR5 positivity on HRS cells. CXCL13+ macrophages surround CXCR5+ HRS cells and PD1+ CD4+ T cells were also present in the tumor-microenvironment. In contrast, CXCR5+ B cells were enriched in r/r CHL with good prognosis.

## Acknowledgements

This study is supported by Program Project Grant funding from the Terry Fox Research Institute (Grant No. 1061), Large Scale Applied Research Project funding from Genome Canada (Grant No. 13124), Genome BC (Grant No. 271LYM) and CIHR (Grant No. GP1-155873), the Canadian Cancer Society Research Institute (Grant No. 705288), a Foundation grant from CIHR (Grant No. 148393), the BC Cancer Foundation and the Paul G. Allen Frontiers Group (Distinguished Investigator award to C.S. Grant No. 12829). T.A. was supported by fellowships from the Japanese Society for The Promotion of Science, the Uehara Memorial Foundation, CIHR and the Lymphoma Research Foundation. T.A. received research funding support from The Kanae Foundation for the Promotion of Medical Science. T.A. recieved research support as a Lymphoma Research Foundation Lymphoma Scientific Research Mentoring Program Scholar.

