## Supplemental File for "The spatially resolved tumor microenvironment predicts treatment outcome in relapsed/refractory Hodgkin lymphoma"

### Supplemental Methods

#### *Tissue samples*

The study cohort and Patient characteristics are summarized in **Appendix Fig 1** and **Appendix Table S1**. Detailed information on the discovery cohort is also described in the previous publication[12]. We analyzed IMC data from 165 CHL samples, including 71 patients with paired primary relapse specimens and 22 diagnostic control samples without any relapse, termed the 'discovery cohort' (**Appendix Fig A1)**[12]. Biopsies of these CHL cases, treated at BC Cancer between 1985 and 2011, are part of a tissue microarray (TMA) that was previously reported[12]. In brief, the patients were selected according to the following criteria: patients received first-line treatment with doxorubicin, bleomycin, vinblastine, and dacarbazine (ABVD) or ABVD-equivalent therapy with curative intent; patients experienced CHL progression after primary treatment (refractory disease or relapse); and tissue derived from an excisional biopsy was available[12]. Patients were classified as having early relapse disease if their CHL progressed within 12 months after initial diagnosis or refractory to first-line treatment. We also assembled a second, independent cohort for validation, termed the ‘validation cohort’ (n = 44) used for Multi-color immunofluorescence (MC-IF). For the validation cohort, we selected r/r CHL patients treated at BC Cancer between 2012 to 2021 with available FFPET relapse biopsies, according to the same selection criteria as for the discovery cohort. For single cell RNA sequencing, three patients with histologically confirmed diagnostic CHL were included in this study. Patients were selected based on the availability of tissue that had been mechanically dissociated and cryopreserved as cell suspensions following diagnostic lymph node biopsy at BC Cancer.

***Tissue microarray (TMA) construction, single color IHC and EBER-1 ISH on TMA***

For TMA construction in the validation cohort, 1.5mm duplicate cores were obtained from representative areas containing Hodgkin Reed-Sternberg cells of 28 relapse biopsies of CHL. The diagnosis was made according to the WHO classification and reviewed by hematopathologists (KT, TT and PF)[45]. For immunohistochemistry (IHC) staining, 4µm slides of the TMA and antibodies listed in **Appendix Table A3** were used. Staining was performed on a Benchmark XT platform (Roche Diagnostics, USA) or intelliPATH platform (Biocare Medical, USA). The slides were independently reviewed and scored by KT and/or TT and/or PF and the single stain IHC scores were utilized for MC-IHC segmentation. Epstein-Barr virus-encoded small RNA 1 (EBER-1) in situ hybridization (ISH) was performed according to the manufacturer’s protocol (Roche Diagnostics, USA).

***Imaging mass cytometry***

Imaging mass cytometry (IMC) was performed on a 5μm section of the same TMA described above. The section was baked at 60°C for 90 minutes on a hot plate, de-waxed for 20 minutes in xylene and rehydrated in a graded series of alcohol (100%, 95%, 80% and 70%) for 5 minutes each. Heat-induced antigen retrieval was conducted using a Sous-Vide cooker at 95°C in Tris-EDTA buffer at pH 9 for 30 minutes. After blocking with 3% BSA in PBS for 45 minutes, the section was incubated overnight at 4°C with a cocktail of 35 antibodies tagged with rare lanthanide isotopes (**Appendix Table A2**). The section was counterstained the next day for 40 minutes with iridium (Ir) nuclear stain[46]. Slides were imaged using the Fluidigm Hyperion IMC system with a 1μm laser ablation spot size and frequency of 200Hz. Tissue areas of approximately 1 $mm^2$ per sample were ablated and imaged. Duplicate cores of the same samples were ablated when morphologic heterogeneity was identified a priori on H&E. Image analyses were performed using CellProfiler (v4.1.3), Ilastik (v1.3.3) and HistoCAT (v1.75). To perform IMC spatial analysis, we selected specific cell types based on marker expression, the number of nearest neighbors, and a spatial interaction range. The spatial interaction range is the distance within which cells are likely to interact, and we chose a range of 50 microns. For each cell, we calculated the spatial interaction score, called 'spatial score', to a given cell type as the distance to the 5 nearest neighbor cells, capped at the spatial interaction range, scaled, and inverted. To ensure the reliability and comparability of spatial scores at the sample level, we calculated the mean of the top 10% of spatial scores of each variable for each sample. By employing this approach, we minimized the bias that zero values of the spatial scores might cause for subsequent analysis at the sample level.

***Prognostic Model Development***

To develop a prognostic model for r/r CHL patients, we applied our new cross-format LASSO_plus algorithm (https://github.com/ajiangsfu/csmpv) on the combination of standardized spatial scores and traditional protein percentages. LASSO_plus, which draws upon the principles of the well-established LASSO method and incorporates single and stepwise variable selection techniques, enabled us to select variables from two separate lists (58 standardized spatial score variables and 61 standardized protein variables). We initially set the top N = 10 for each list, then combined them and ran a Cox model on the resulting combined list. We retained variables that were selected from only one data format, and for duplicated variables from both formats, we only kept those with p-values <= 0.05. As a result, we obtained a list of six variables (**Figure 2A**). To make the model more practical, we manually removed two variables whose end of hazard ratios' 95% CI is overlapped with the range of 0.95-1.05 (GzMB pDC spatial score or TIM3 Treg spatial score). This left us with four variables: CXCR5 HRS spatial score, PD1 CD4 spatial score, Mac (macrophage) spatial score, and CXCR5 B spatial score. We chose to use only the spatial score data format. We used our new algorithm Xgpred in csmpv R package (https://github.com/ajiangsfu/csmpv) to build the model, which first built an XGBoost[47] (eXtreme Gradient Boosting) model on the four selected variables for post-ASCT outcome. To achieve stable high and low-risk groups based on the XGBoost model score, we applied spline regression, model-based clustering techniques, and additional filtering steps. After obtaining stable high and low-risk samples, we used known classifications to build a linear regression for each of the four variables to obtain t-values, which were treated as variable weights together with the four variables to build the predictive model with a Linear Prediction Score

(LPS), called RHL4S. Finally, we utilized an empirical Bayesian approach to calculate the probability of a sample being classified as a high-risk sample. Based on a probability cut-off of 0.8, any sample with a probability of being in a high-risk group of >= 0.8 was classified as high-risk, and the remaining samples were classified as low-risk.

***Multi-color immunofluorescence (MC-IF) on TMA, scanning and image analysis***

TMA slides were deparaffinized in xylene and rinsed with $dH_2O$. Antigen retrieval was performed in AR6 buffer (PerkinElmer, USA) with Diva decloaker (Biocare Medical, USA). The primary antibody for CXCL13 was incubated for 30min in an Intellipath FLX rack at room temperature, followed by detection using the Mach2 mouse HRP with 10 min incubation. Visualization of CXCL13 was achieved using Opal 520. The slide was placed into AR6 buffer and heated using a microwave. In serial order, the slide was incubated with primary antibody for CXCR5, followed by detection using Mach2 rabbit HRP, and visualization was accomplished using Opal 650. The slide was again placed into AR6 buffer and heated using a microwave. Then the primary antibody for CD68 was incubated, followed by detection of Mach2 rabbit HRP and Opal 550 for visualization. The slide was placed into AR6 buffer for microwaving. The primary antibody for CD4 was incubated, followed by detection of Mach2 rabbit HRP and visualization for Opal 650. Microwave heating was repeated again with AR6 buffer. The primary antibody for PD-1 was incubated for 30min in an Intellipath FLX rack at room temperature, followed by detection using the Mach2 mouse HRP with 10 min incubation. Visualization of PD-1 was achieved using Opal 620. The primary antibody for CD30 was incubated, followed by detection of Mach2 mouse HRP and visualization for Opal 570. Nuclei were visualized with DAPI staining and the section was coverslipped using Fluoro Care Anti-Fade Mountant. TMA slides were scanned using the Vectra multispectral imaging system (PerkinElmer, USA) following manufacturer's instructions to generate .im3 image cubes for downstream analysis. Optimal exposure times for fluorophores ranged between 50 and 200ms. To analyze the spectra for all fluorophores included, inForm image analysis software (v2.4.4; PerkinElmer, USA) was used. Cells were first classified into tissue categories using DAPI and CD30 to identify $CD30^{+}DAPI^{+}$, $CD30^{-}DAPI^{+}$, and $CD30^{-}DAPI^{-}$ areas via manual circling and training[13]. The $CD30^{+}DAPI^{+}$ regions were considered to be HRS-surrounding regions. Cells were then phenotyped as positive or negative for each of the six markers (CXCL13, CXCR5, CD68, CD4, CD30, PD-1) or (CD20, CXCR5, CD68, CD4, CD30, PD-1). Data were merged in R by X-Y coordinates so that each cell could be assessed for all markers simultaneously. Nearest neighbor analysis was performed with the spatstat R package (v1.58-2).

#### *Survival analysis*

Overall survival (OS) was defined as the time from diagnosis to death from any cause. Time to first relapse was defined as the time from primary diagnosis to first CHL progression, or death from CHL. Post-ASCT-OS was defined as time from ASCT

treatment to death from any cause. Post-ASCT-FFS was defined as time from ASCT treatment to CHL progression/relapse, or death from any cause. Patients with complete or partial response after second-line chemotherapy were classified as chemo-sensitive. Patients with stable or progressive disease were classified as chemo-resistant. Non-parametric survival analyses with a single binary predictor were analyzed using the Kaplan-Meier method and results were compared using the log rank test. Univariate and multivariate Cox regression analyses were performed to assess the effects of prognostic factors. Survival analyses were performed in the R statistical environment (v4.2.2).

***Prognostic Model Development***

To develop a prognostic model for r/r CHL patients, we applied our new cross-format LASSO_plus algorithm (https://github.com/ajiangsfu/csmpv) on the combination of standardized spatial scores and traditional protein percentages. LASSO_plus, which draws upon the principles of the well-established LASSO method and incorporates single and stepwise variable selection techniques, enabled us to select variables from two separate lists (58 standardized spatial score variables and 61 standardized protein variables). We initially set the top N = 10 for each list, then combined them and ran a Cox model on the resulting combined list. We retained variables that were selected from only one data format, and for duplicated variables from both formats, we only kept those with p-values <= 0.05. As a result, we obtained a list of six variables (Figure 2A). To make the model more practical, we manually removed two variables whose hazard ratios' 95% CI is overlapped with the range of 0.95-1.05 (GzMB pDC spatial score or TIM3 Treg spatial score). This left us with four variables: CXCR5 HRS spatial score, PD1 CD4 spatial score, Mac (macrophage) spatial score, and CXCR5 B spatial score. We chose to use only the spatial score data format, as it was the dominant data format and CXCR5 B is the only variable out of four variables that is in protein percentage format, resulting in our final variable list for the predictive model. We used our new algorithm XGpred in csmpv R package (https://github.com/ajiangsfu/csmpv) to build the model, which first built an

XGBoost[47] (eXtreme Gradient Boosting) model on the four selected variables for post-ASCT outcome. To achieve stable high and low-risk groups based on the XGBoost model score, we applied spline regression, model-based clustering techniques, and additional filtering steps. After obtaining stable high and low-risk samples, we used known classifications to build a linear regression for each of the four variables to obtain t-values, which were treated as variable weights together with the four variables to build the predictive model with a Linear Prediction Score (LPS), called RHL4S. Finally, we utilized an empirical Bayesian approach to calculate the probability of a sample being classified as a high-risk sample. Based on a probability cut-off of 0.8, any sample with a probability of being in a high-risk group of >= 0.8 was classified as high-risk, and the remaining samples were classified as low-risk. XGpred.predict function in the same package can be used for prediction on a new data set when comparable data is available.

***RHL4S packages***

Since our ultimate goal is to establish prognostic model, which can be readily generalized and applicable to patients in daily clinical practice, we translated the findings from IMC into more simplified data. For this aim, we developed the RHL4S R package (https://github.com/ajiangsfu/RHL4S). The RHL4S calls function is a wrap-up function designed to process MC-IF data, calculate and calibrate the RHL4S model scores, and classifies patients as either high or low risk for each patient.

***Single cell RNA sequencing sample preparation***

To purify HRS cells, we implemented a flow cytometry-based cell sorting approach based on published literature[17,18]. Cell suspensions from CHL tumors were rapidly defrosted at 37°C, washed in RPMI1640/20% FBS solution containing Dnase I (Millipore Sigma, Darmstadt, Germany) and washed in PBS containing 2% FBS. Cells were resuspended in PBC containing 2% FBS and stained with antibody panel specific to isolate HRS cells (**Appendix TableA4**) for 15 minutes at 4°C in the dark. Viable cells (DAPI negative) were sorted on a FACS ARIAIII or FACS Fusion (BD Biosciences) using a 130 µm nozzle (**Appendix FigA7**) and were analyzed using FlowJo software (v10.2; TreeStar, Ashland, OR, USA). Sorted cells were collected in 0.3 mL of medium, centrifuged and diluted in 1x PBS with 0.04% bovine serum albumin (BSA). Cell number was determined using a Countess II Automated Cell Counter whenever possible.

***Hybrid capture sequencing of marker genes using CapID***

Hybrid capture sequencing was performed as described previously[19]. In brief, Probes for 177 genes (**Appendix Table A5**) were designed and synthesized by Twist Bioscience. Hybridization capture of DNA libraries was performed using Twist Hybridization and Wash Kit (Twist Bioscience). First, we pooled 500 ng of each library
to multiplex 8 libraries in a low-bind tube and performed hybridization according to the Twist Target Enrichment Protocol. The captured library was measured using Agilent Bioanalyzer High Sensitivity chip and Qubit dsDNA HS Assay Kit and run on Illumina Nextseq550.

***Normalization and batch correction***

Analysis and visualization of scRNA-seq data was performed in the R statistical environment (v4.1.0). CellRanger software (v6.0.2) was used to align the sequencing reads to the hg38 human reference genome build. CellRanger (v6.0.2) count data from all cells (n = 3302) were read into a single 'Seurat' object using the Seurat Package (v4.2.1)[48]. Cells were filtered if they had ≥ 20% reads aligning to mitochondrial genes, or if their total number of feature counts was less than 1000. This yielded a total of 3302 cells for analysis. The read count matrix was used as input into the "NormalizeData" function which returned a normalized expression matrix. Principal component analysis (PCA) was then run on the normalized expression matrix using highly variable genes identified by "FindVariableGenes" function.

#### *Clustering and annotation*

Unsupervised clustering was performed with the "FindClusters" function[48], using the first 30 PCA components as input. Clusters were manually assigned to a cell type by comparing the mean expression of known markers across cells in a cluster. Markers used to annotate cells included CD19 (B cells), CD8, CD3, CD4 (T cells), CD68 (Macrophages) and CCL17, TNFRSF8 (HRS cells). The clustering results were shown in UMAP space which was generated using the first 30 PCA components.

#### *Cell to cell interaction analysis*

**Cell Chat analysis.** The CellChat R package (v1.1.3)[31] was used to identify potential cell-cell communication networks from scRNA-seq data. Cells were classified into broad subtypes based on their cluster assignments (i.e. HRS-C1, HRS-C2 and etc.) which were input into CellChat as cell labels. The ligand-receptor interaction database (n = 1999) used for predicting intercellular communications was the CellChat built-in cross-referencing ligand-receptor interaction database (n = 1939) and manually curated ligand-receptor interactions (n = 60) which were important in CHL TME biology. The communication probability was calculated for each ligand-receptor pair and the significant interactions were identified (P-values < 0.05). The CellChat results were visualized using the "LRPlot" from the iTALK R package (v0.1.0)[32].

***RHL30 Prognostic Model/Assay***

RHL30 scores were calculated using methods described in Chan et al[12]. In brief, RNA extracted from formalin-fixed paraffin-embedded tissue (FFPET) were hybridized to RHL30 CodeSet for 12–30 h at 65°C. The RHL30 is a 30 probe NanoString codeset comprising of 18 endogenous genes and 12 housekeepers. The samples were then run on an nCounter Digital Analyzer (Nanostring, Seattle, WA, USA). Then, quality control was performed, and gene expression data were normalized and RHL30 scores were calculated using the same methods as described before.  We used the median as a cut-off to distinguish cases with low and high-risk since the original cut-off was not suitable due to batch effects.

***Statistical results & visualization***

All t-tests reported are two-sided Student's t-tests, and P-values < 0.05 were considered statistically significant. In all boxplots, boxes represent the interquartile range with a horizontal line indicating the median value. Whiskers extend to the farthest data point within a maximum of 1.5 × the interquartile range, and colored dots represent outliers.

***Data availability***

Single cell RNA-seq counts (generated with CellRanger v2.1.0) and a merged 'SingleCellExperiment' R object is available in the European Genome-phenome Archive (EGA) (TBA) via controlled access.

***Code availability***

Scripts used for data analysis are available upon request. Our in-house R packages, csmpv (https://github.com/ajiangsfu/csmpv) and RHL4S (https://github.com/ajiangsfu/RHL4S) are available on GitHub.

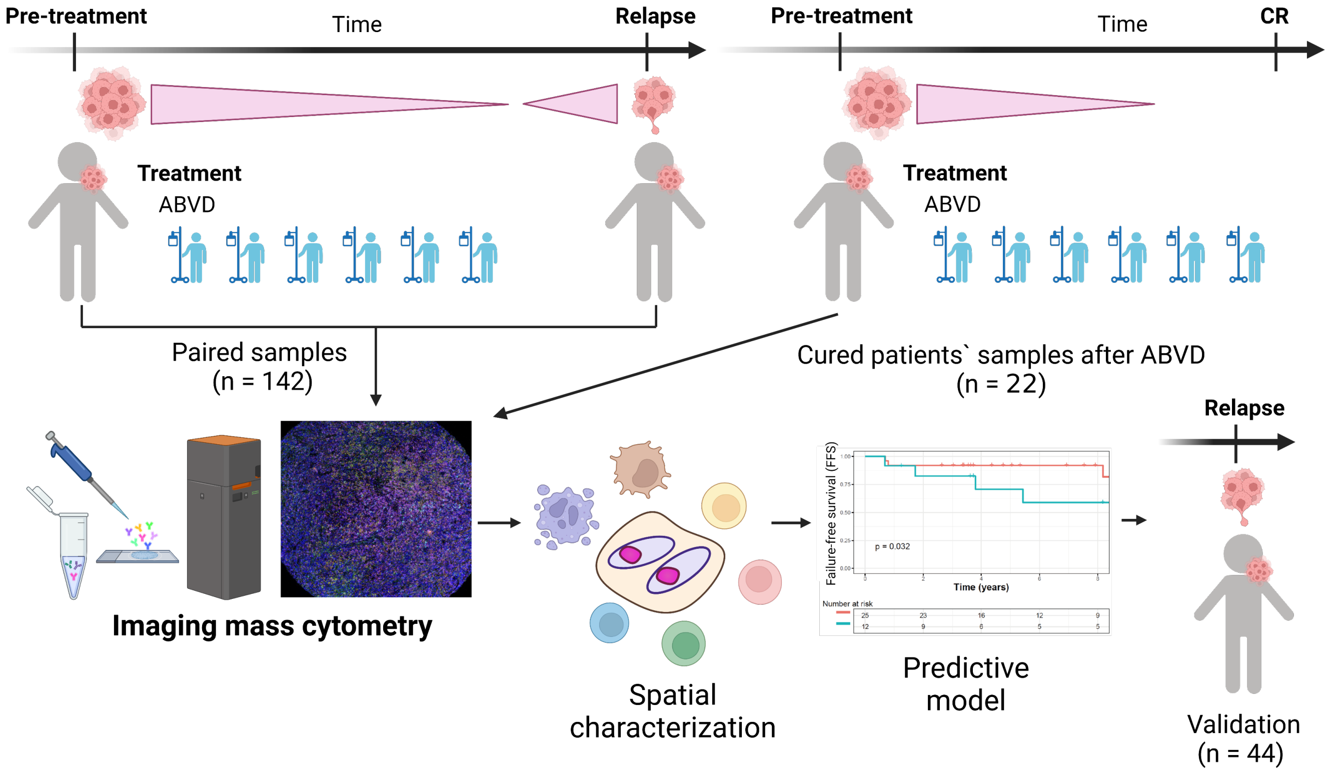


**Appendix FigA1. Cohort and study design Overview.** We analyzed IMC data from 164 CHL samples, including 71 patients with paired primary relapse specimens and 22 diagnostic control samples without any relapse. Subsequently, we developed a novel predictive model using spatial information.The predictive model was validated using an independent validation cohort of 44 patients.

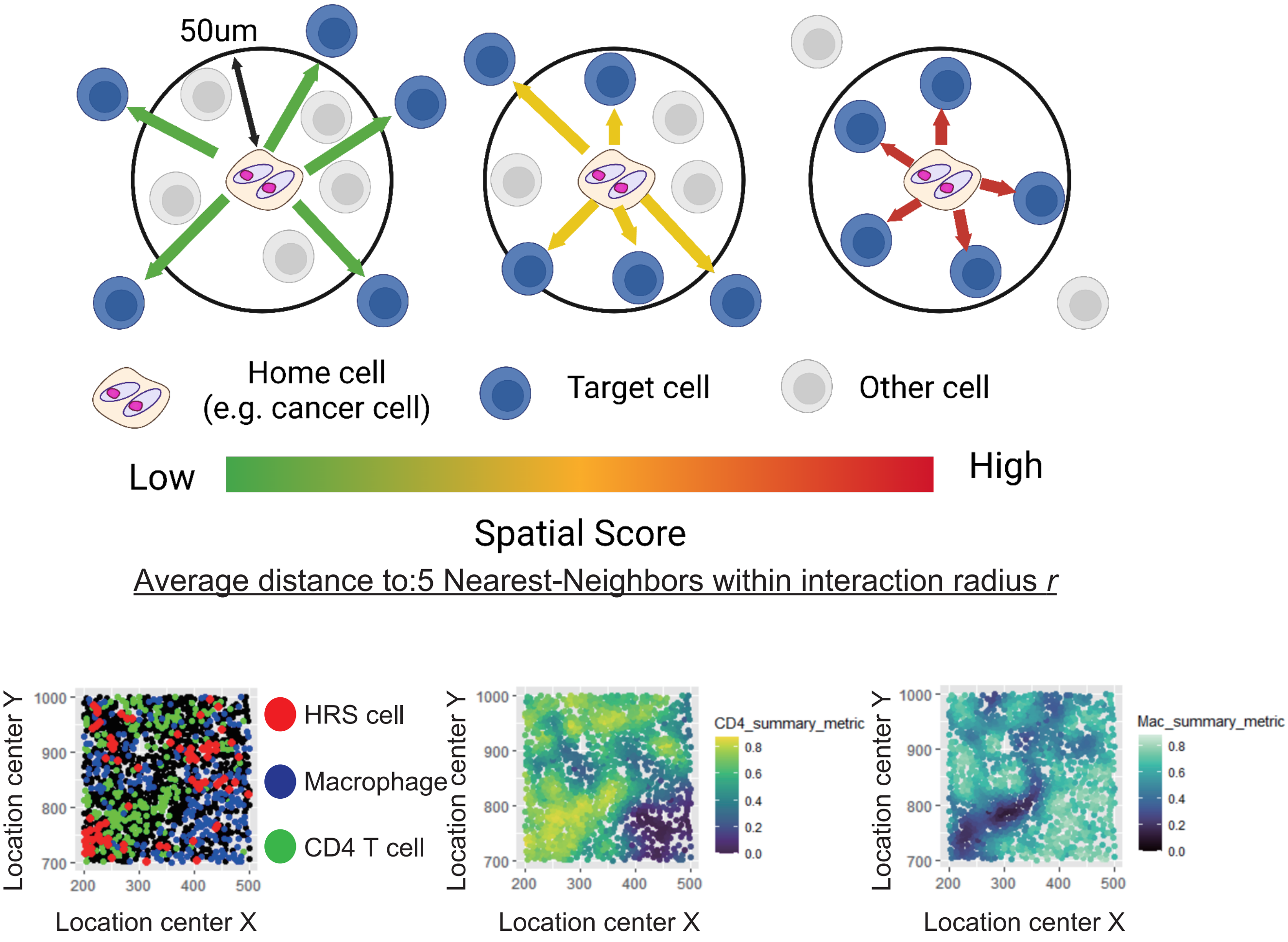


**Appendix FigA2. Scheme of spatial score.** Average distance to 5 nearest neighbors within interaction radius r from cells of interests (home cells) is calculated and scored. Representative images of regions with CD4 T cell (CD4) enrichment (center, green) and macrophage (Mac, blue) abundance (right) with HRS cell (Red) are shown at the bottom.

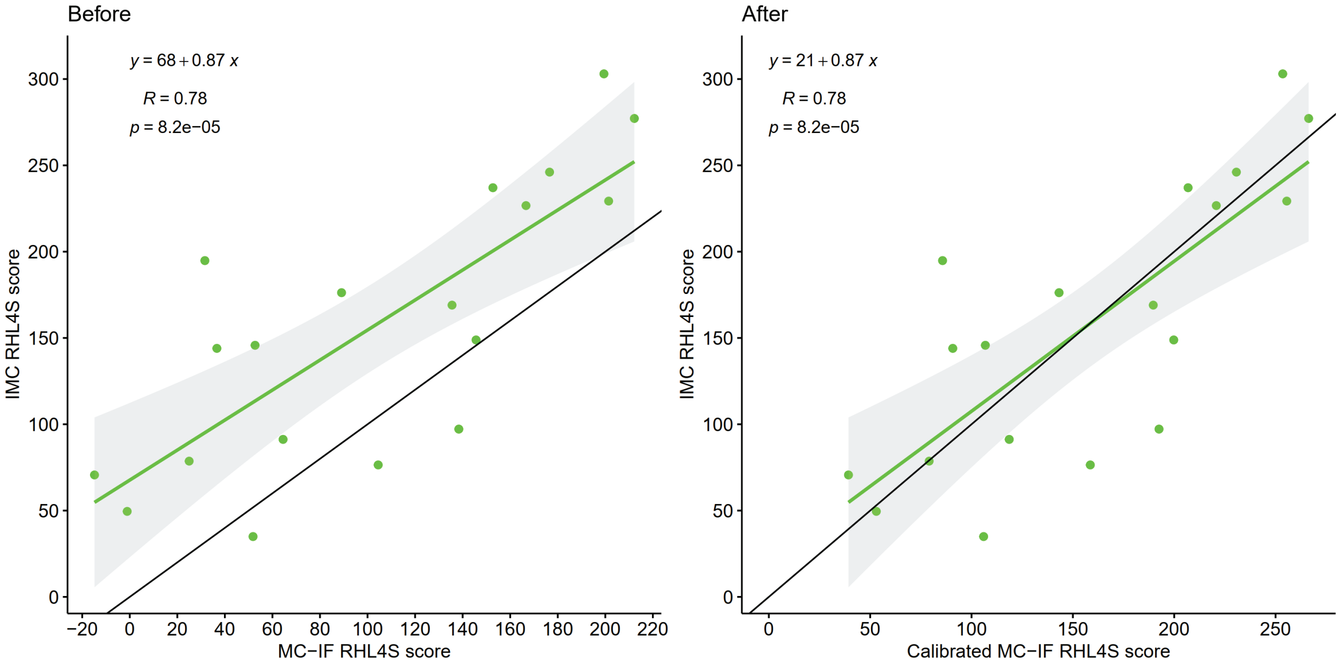


**Appendix Fig A3. Calibration plots for RHL4S from imaging mass cytometry (IMC) and multi-color immunofluorescence (MC-IF).** Plots show an association between RHL4S from IMC and RHL4S from MC-IF before (left) and after (right) calibration.

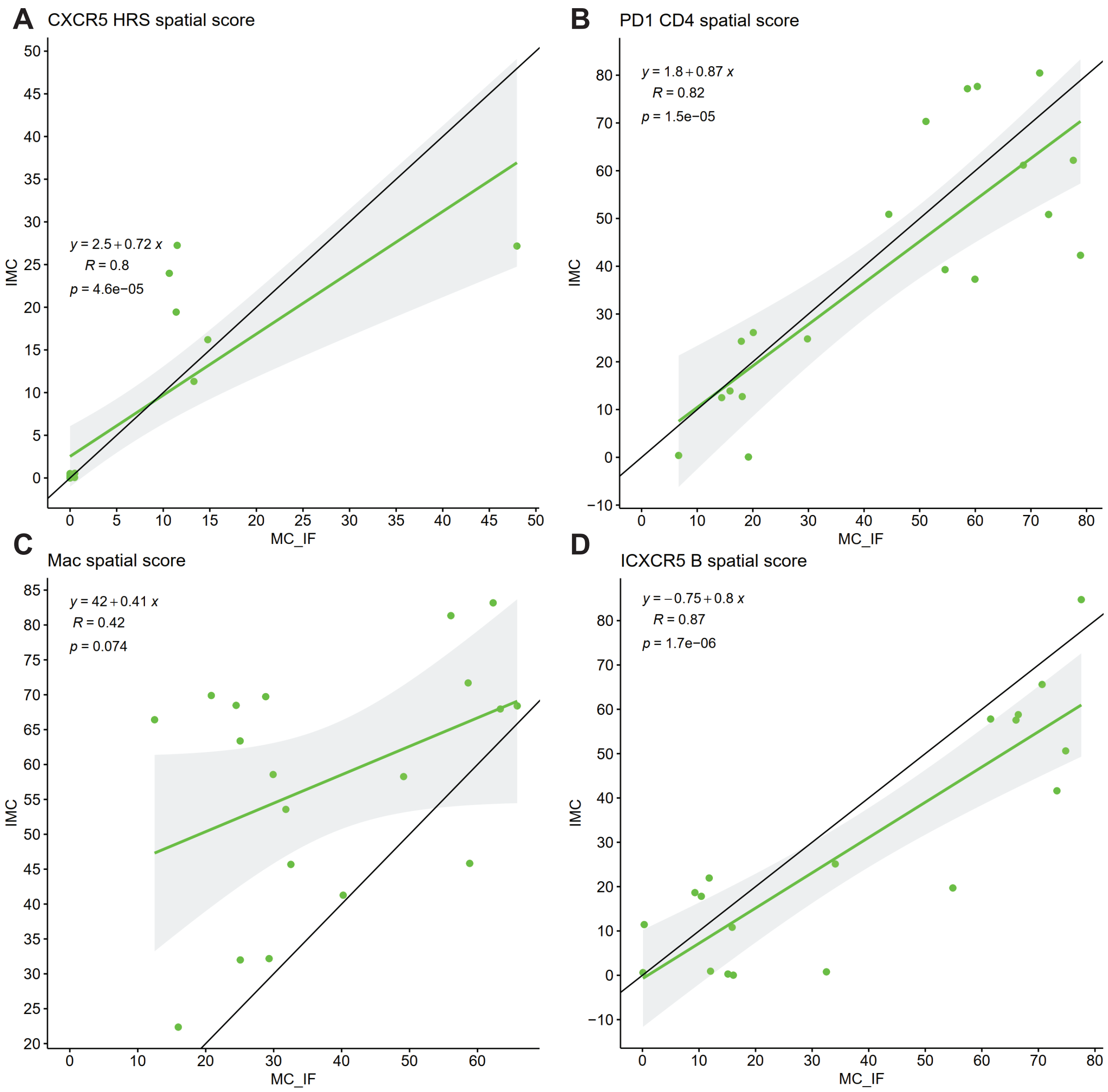


**Appendix Fig A4. Correlation plots of spatial scores of 4 variables for RHL4S.** The plot shows the association between spatial scores of 4 variables for RHL4S between data from imaging mass cytometry and data from multi-color immunofluorescence.

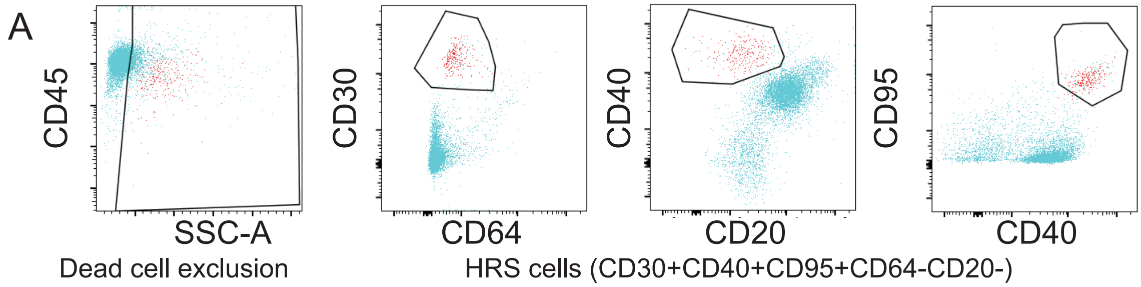


**Appendix Fig A5. Gating strategy used to sort HRS cells in cell suspensions from classic Hodgkin lymphoma.**

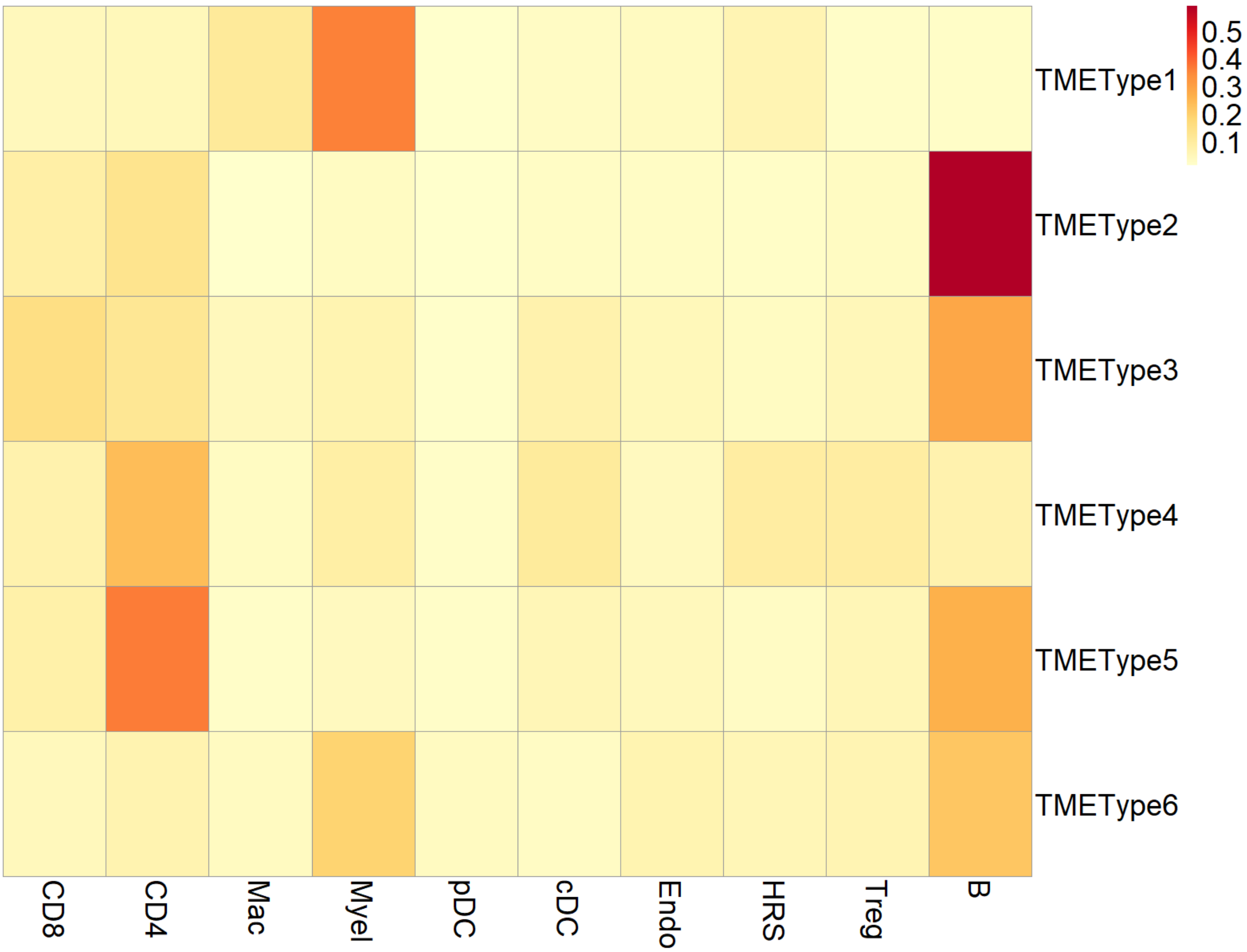


**Appendix FigA6. Tumor-microenvironment subtype.** Heatmap summarizing the enrichment of selected immune cell subtypes and HRS cells clustered using the Phenograph algorithm defined by Imaging Mass Cytometry data. Six tumor-microenvironment subtypes were identified.

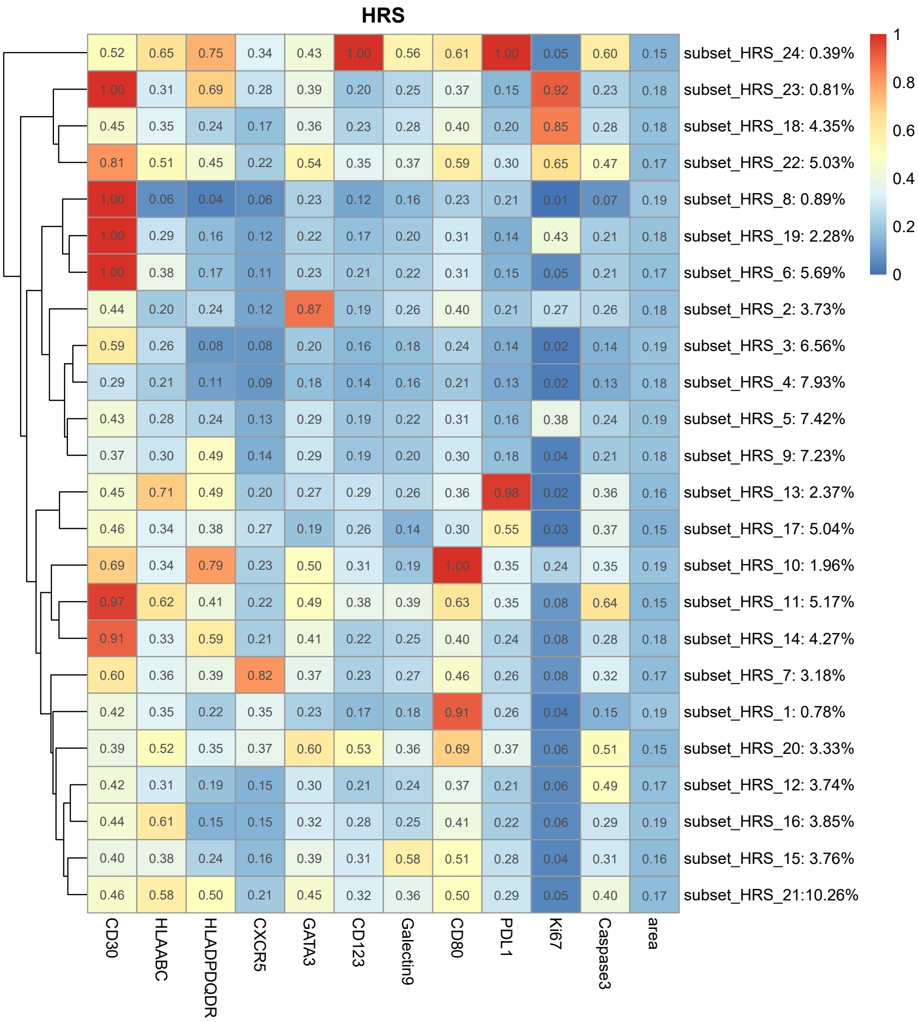


**Appendix FigA7. Subset of HRS cells.** Heatmap summarizing the median expression of selected protein markers on HRS cells clustered using Phenograph. Prior to clustering protein expression values were Arcsinh transformed with a cofactor of 5, clipped at the 99th percentile, and scaled from 0 to 1. The dendrogram represents hierarchical clustering of the heatmap rows (HRS subset clusters) based on Euclidean distance.

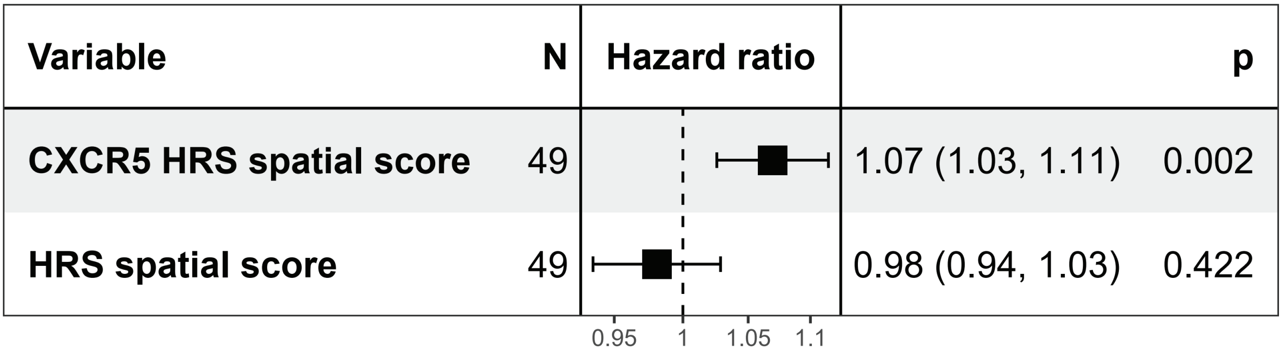


**Appendix Fig A8. CXCR5+ HRS cells vs. HRS cells Forest Plot.** The spatial score of the CXCR5+ HRS cells subset is compared to the spatial score of all HRS cells of post-ASCT FFS as the endpoint in pairwise Cox regression. Each dot represents the hazard ratio (x-axis) with the bar representing the 95% confidence interval.

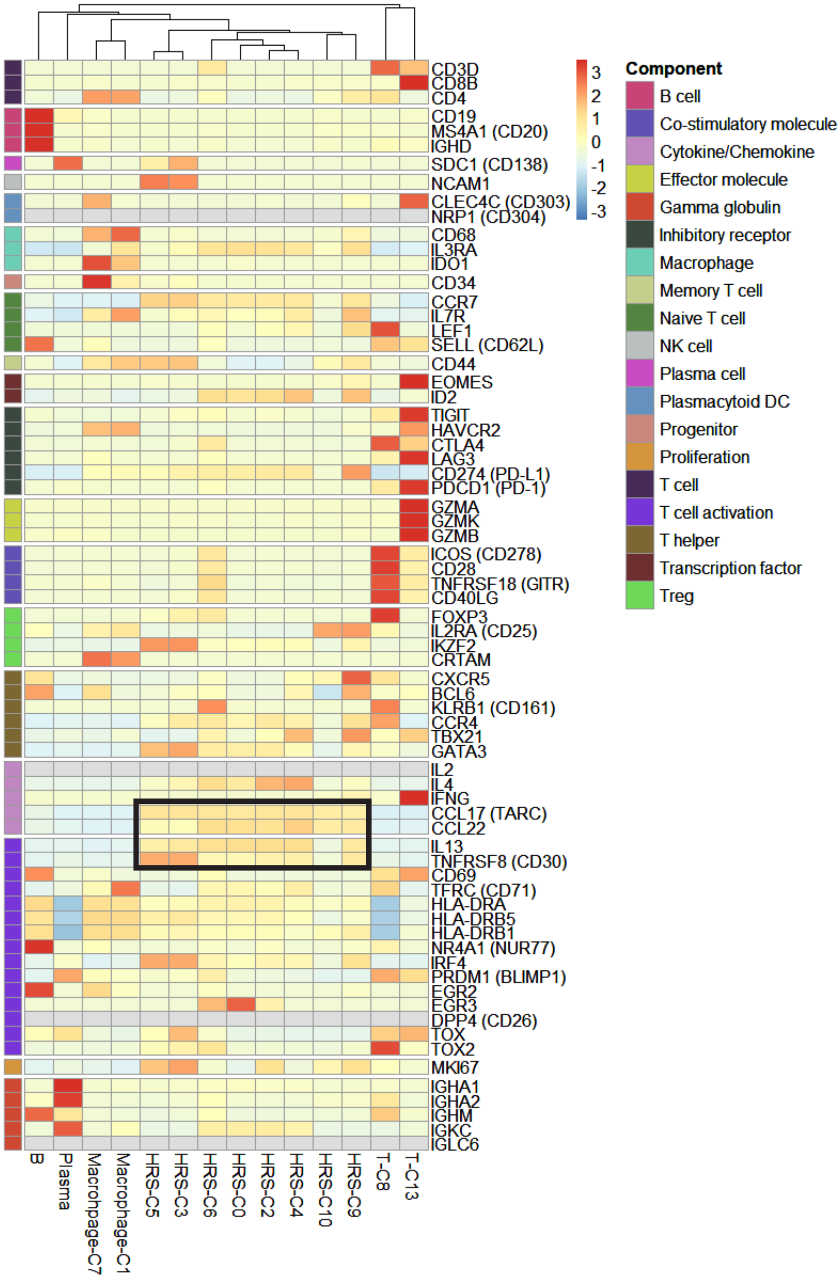


**Appendix Fig A9. Single cell RNA sequencing of enriched HRS cells.** Heatmap summarizing mean expression (normalized and log transformed) of selected canonical markers in each cluster. Data have been scaled row-wise for visualization. The covariate bar on the left side indicates the component associated with each gene, and black boxes highlight prominent expression of known HRS cell genes.

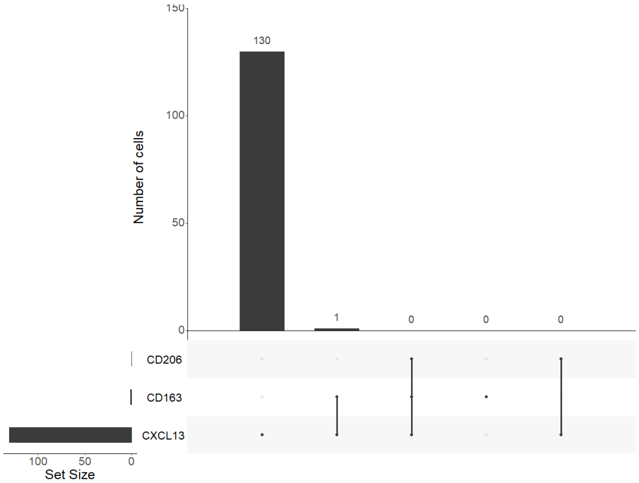


**Appendix Fig A10. Co-expression pattern of CXCL13+ macrophages.** UpSet plot showing co-expression patterns of M2 macrophage markers (CD163 and CD206) in CXCL13+ macrophages by single cell RNA sequencing.

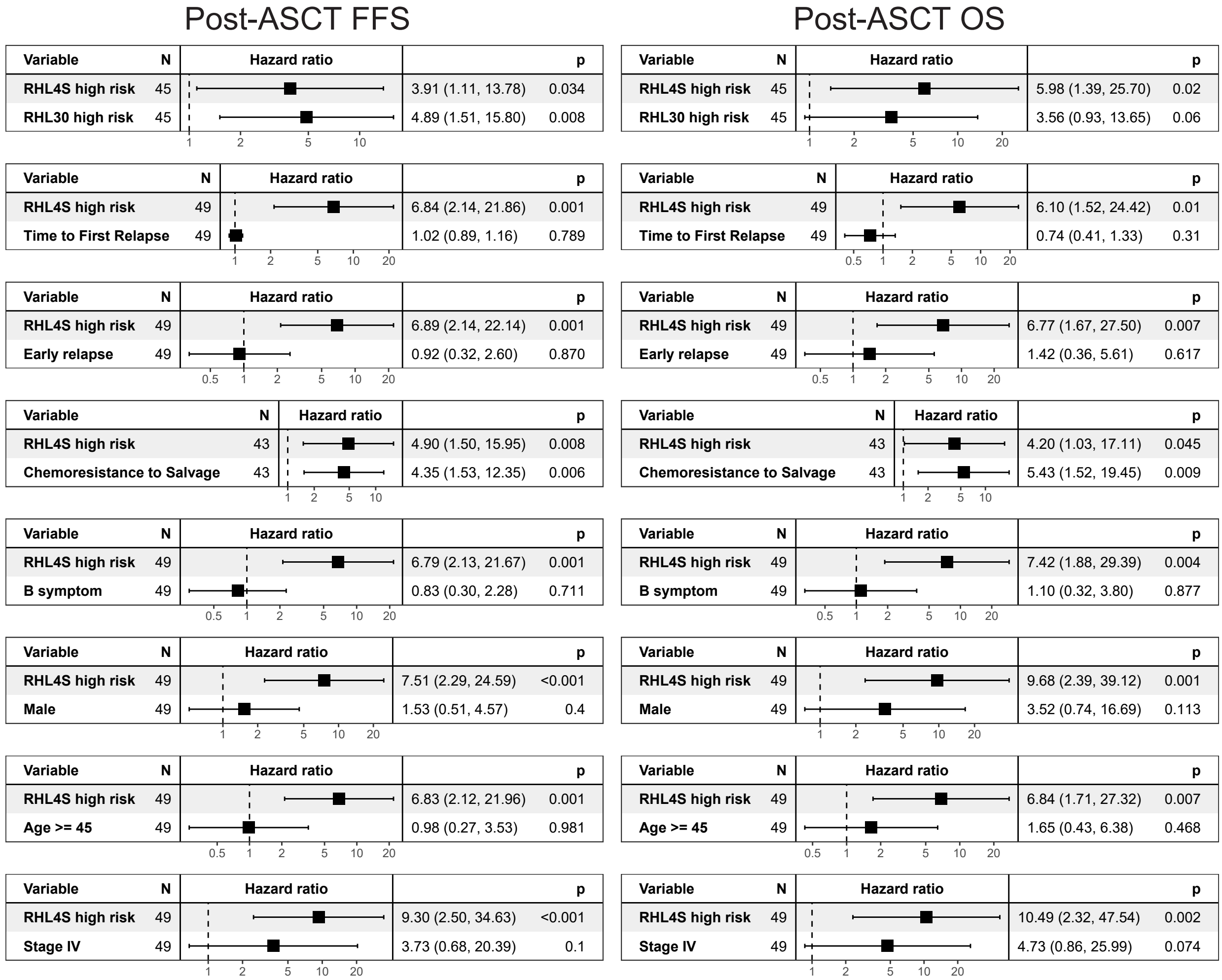


**Appendix Fig A11. RHL4S vs. Reported Prognostic Markers Forest Plot.** The RHL4S risk-class is compared to other reported prognostic markers (y-axis) of (Left) post-ASCT failure-free survival (FFS) and (Right) post-ASCT overall survival (OS) using pairwise Cox regression of two variables. Each dot represents the hazard ratio (x-axis) with the bar representing the 95% confidence interval. Each facet on the y-axis is a different pairwise multivariate Cox regression.

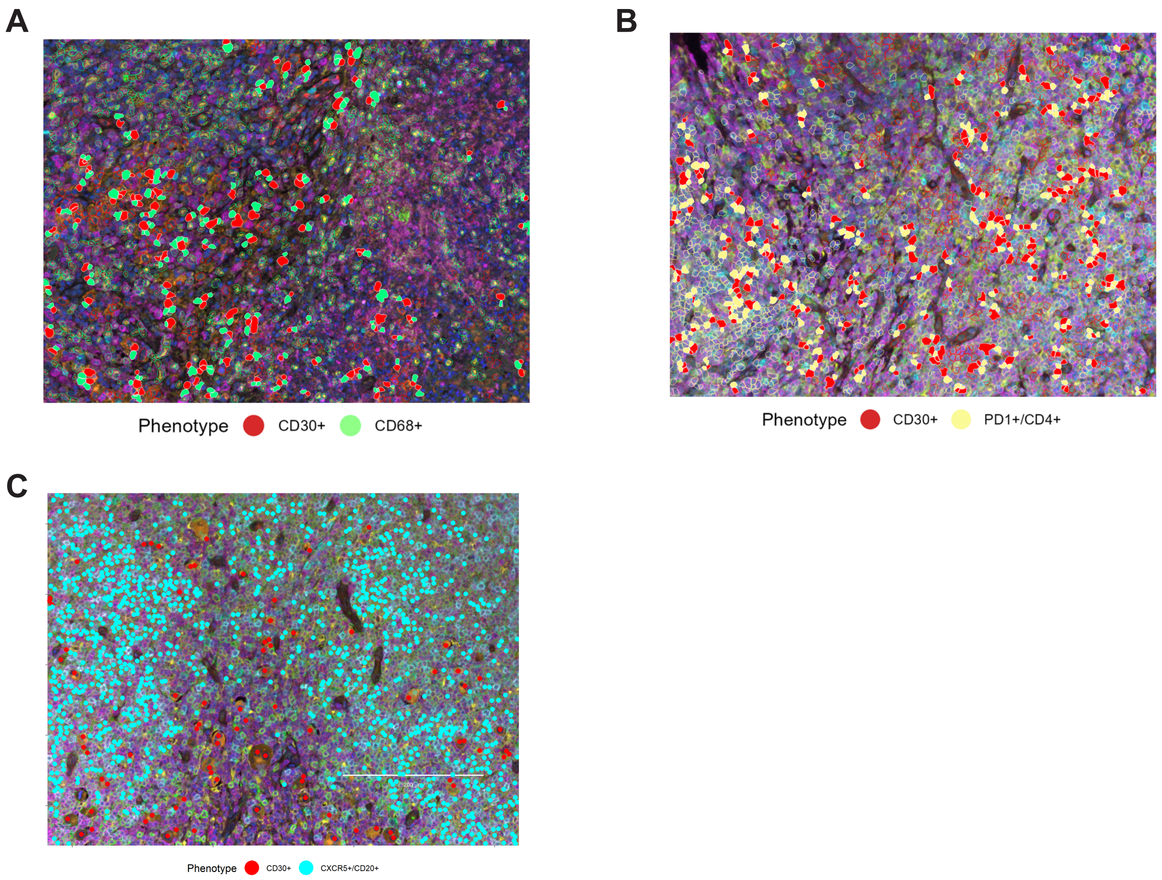


**Appendix Fig A12. Multi-color immunohistochemistry in the validation cohort.** Membrane map depicting (A) CD68+ Macrophage (green) and CD30+ HRS cells (red). (B) CD4+ PD1+ T cell (yellow) and (C) CXCR5+ B cells.

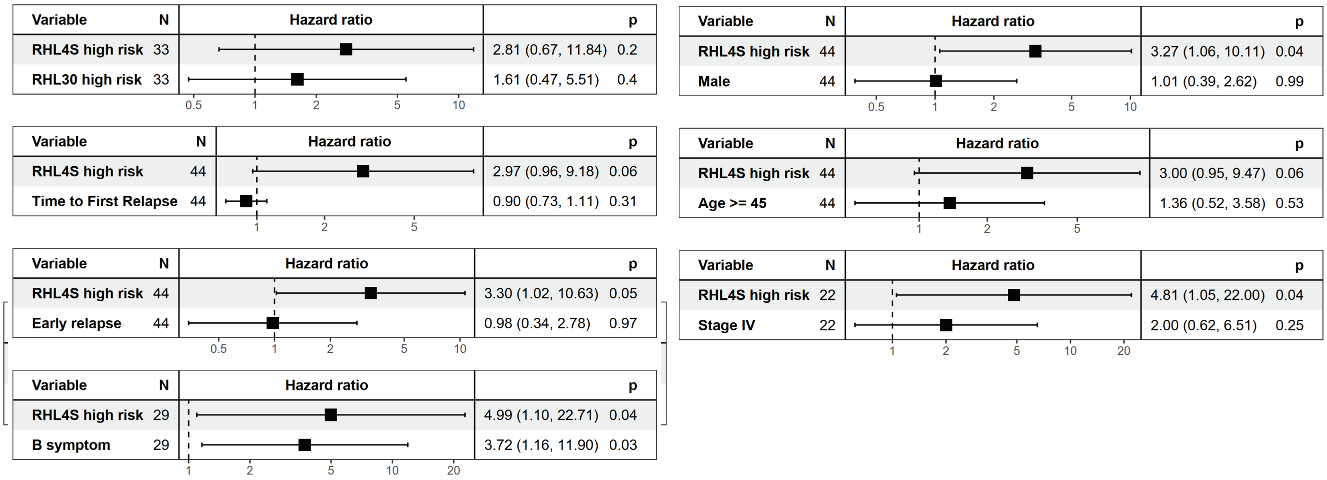


**Appendix Fig A13. RHL4S vs. Reported Prognostic Markers Forest Plot in validation cohort.** The RHL4S risk-class is compared to other reported prognostic markers (y-axis) of post-ASCT outcomes using pairwise Cox regression of two variables. Each dot represents the hazard ratio (x-axis) with the bar representing the 95% confidence interval. Each facet on the y-axis is a different pairwise multivariate Cox regression.

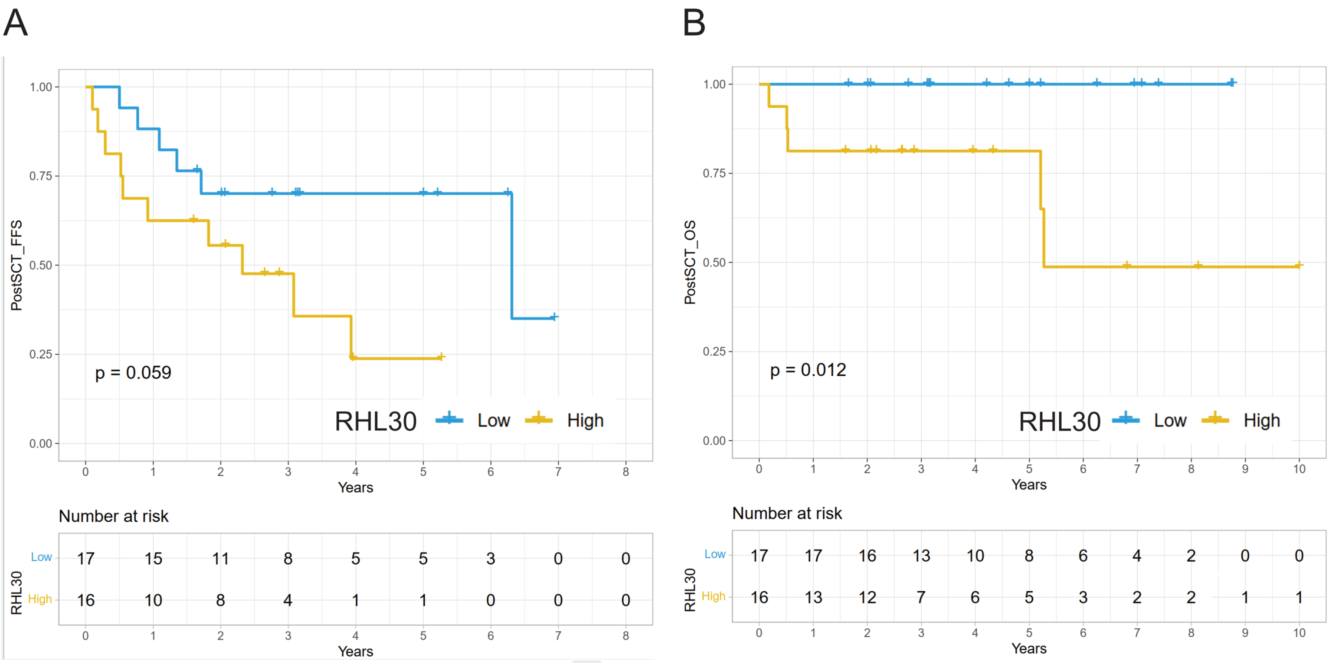


**Appendix Fig A14. Post-ASCT failure-free survival according to risk group of RHL30 in validation cohort.** Kaplan-Meier curves of the high- versus low-risk groups for (A) post-ASCT failure-free survival (FFS) and (B) post-ASCT overall survival (OS) as identified by RHL30 in independent validation cohort, respectively. P values were calculated using a log rank test.
